# A quantitative, genome-wide analysis in *Drosophila* reveals transposable elements’ influence on gene expression is species-specific

**DOI:** 10.1101/2022.01.20.477049

**Authors:** Marie Fablet, Judit Salces-Ortiz, Angelo Jacquet, Bianca F. Menezes, Corentin Dechaud, Philippe Veber, Rita Rebollo, Cristina Vieira

**Affiliations:** Laboratoire de Biométrie et Biologie Evolutive, Université de Lyon; Université Lyon 1; CNRS; UMR 5558, Villeurbanne, France; Institut Universitaire de France (IUF); Institut de Génomique Fonctionnelle de Lyon, Univ Lyon, CNRS UMR 5242, Ecole Normale Supérieure de Lyon, Université Claude Bernard Lyon 1, 46 allée d’Italie, F-69364 Lyon, France; Univ Lyon, INRAE, INSA-Lyon, BF2I, UMR 203, 69621 Villeurbanne, France; Institute of Evolutionary Biology (CSIC-Universitat Pompeu Fabra) P° Marítimo de la Barceloneta, 37-49 08003 Barcelona, Spain; Symbiotron; FR3728 Biodiversité, Eau, Environnement, Ville, Santé; Université Claude Bernard Lyon 1; Villeurbanne 69622, France; Federal Institute of Rio de Janeiro (IFRJ), Pinheiral – RJ, Brazil

**Keywords:** transposon, retrotransposon, transcriptomics, fruit fly, histone

## Abstract

Transposable elements (TEs) are parasite DNA sequences that are able to move and multiply along the chromosomes of all genomes. They can be controlled by the host through the targeting of silencing epigenetic marks, which may affect the chromatin structure of neighboring sequences, including genes. In this study, we used transcriptomic and epigenomic high-throughput data produced from ovarian samples of several *Drosophila melanogaster* and *Drosophila simulans* wild-type strains, in order to finely quantify the influence of TE insertions on gene RNA levels and histone marks (H3K9me3 and H3K4me3). Our results reveal a stronger epigenetic effect of TEs on ortholog genes in *D. simulans* compared to *D. melanogaster*. At the same time, we uncover a larger contribution of TEs to gene H3K9me3 variance within genomes in *D. melanogaster*, which is evidenced by a stronger correlation of TE numbers around genes with the levels of this chromatin mark in *D. melanogaster*. Overall, this work contributes to the understanding of species-specific influence of TEs within genomes. It provides a new light on the considerable natural variability provided by TEs, which may be associated with contrasted adaptive and evolutionary potentials.

**Significance Statement:** Transposable elements (TEs) are parasitic DNA sequences that are widespread components of all genomes. In this study, we combined genomic, transcriptomic and epigenomic high-throughput data produced from ovarian samples of *Drosophila melanogaster* and *Drosophila simulans* wild-type strains, in order to finely quantify the genome-wide influence of TE insertions on gene expression. Our results uncover contrasted patterns depending on the strain, which may have evolutionary impacts.

## Introduction

Transposable elements (TEs) are parasite DNA sequences that are able to move and multiply along the chromosomes of all genomes (Wells & Feschotte 2020). They are source of mutations and genome instability if uncontrolled (Biémont & Vieira 2006; Malone & Hannon 2009; Senti & Brennecke 2010). Control of TEs generally consists in the targeting of particular chromatin marks to TE copies, which induce transcriptional gene silencing and may spread to neighboring sequences and impact gene expression. In this regard, few attempts have been made to finely analyze and quantify TEs’ influence at the whole genome scale (Cridland et al. 2015; Hollister & Gaut 2009; Huang et al. 2016; Lee & Karpen 2017; Uzunović et al. 2019; Wei et al. 2022). In addition, since the very beginning of TE studies, species-specific differences in TE contents, activities and control pathways have been reported in nature, and particularly between *D. melanogaster* and *D. simulans* (Akkouche et al. 2013, 2012; Fablet et al. 2014; Kofler, Nolte, et al. 2015; Lee & Karpen 2017; Mérel et al. 2020; Vieira et al. 2012, 1999). Previous research described the effects of TE insertions on gene expression using collections of strains of *D. melanogaster* (Cridland et al. 2015; Everett et al. 2020; Osada et al. 2017; Zhang et al. 2020), and other studies focusing on a few TE families in wild-type strains of *D. simulans* and *D. melanogaster* uncovered between-species differences in histone mark landscapes (Rebollo, Horard, et al. 2012). Lee and Karpen (Lee & Karpen 2017) provided an analysis on the repressive histone mark H3K9me2 (Histone 3 Lysine 9 dimethylation) around TEs from two *Drosophila* Genetic Reference Panel (DGRP) strains (*D. melanogaster*), and concluded that TE epigenetic effects were pervasive. However, rather than H3K9me2, it is H3K9me3 (Histone 3 Lysine 9 trimethylation) that is known to be associated with the activity of dual-stranded piRNA clusters and the production of TE-derived silencing piRNAs (Le Thomas et al., 2013; Mohn et al., 2014; Sienski et al., 2012). H3K9me3 differs from H3K9me2 in that it is more strongly bound by Rhino, which is abundant in ovaries and leads to piRNA production through alteration of the local transcription program (Mohn et al. 2014).

Several limitations remain from the previous studies, which we propose to address in the present work. First, we connect TE insertion polymorphism, RNA-seq, ChIP-seq on two histone marks (H3K4me3 and H3K9me3), and small RNA-seq data on the same strains. We use eight previously characterized, wild-type strains of *D. melanogaster* and *D. simulans* (Mohamed et al. 2020) that are derived from samples collected in France and Brazil, two strains per location and per species. Using the Oxford Nanopore long read sequencing technology, we previously produced high quality genome assemblies at the chromosome resolution for each strain, which provides us with the various TE insertion sites in each genome (Mohamed et al. 2020). Second, all data are produced from ovaries, *i.e.* the exact same tissue and not mix of tissues. In females, Rhino is known to bind to H3K9me3 and promote the non-canonical transcription of dual-stranded piRNA clusters, in ovaries only (Mohn et al. 2014). Therefore, we expect the strongest control of TEs in this tissue and thus potentially the strongest impact on neighboring genes. In particular, we can speculate that genes located nearby TE insertions may be affected by the local production of piRNAs and hence we searched for gene-derived piRNAs, in association with increased levels of H3K9me3 deposition on gene sequences. We also studied H3K4me3 (Histone 3 Lysine 4 trimethylation), which is known to be associated with active, canonical transcription. Third, the production of genome-wide data from four wild-type strains of *D. melanogaster* and four wild-type strains of *D. simulans* brings the opportunity to statistically test for species-specific differences and provide a quantitative assessment of the contribution of TEs to gene expression, in a comparative genomics perspective (Figure 1). In addition, the use of linear models allows us to finely quantify and compare the contributions at different levels.

**Fig. 1.**
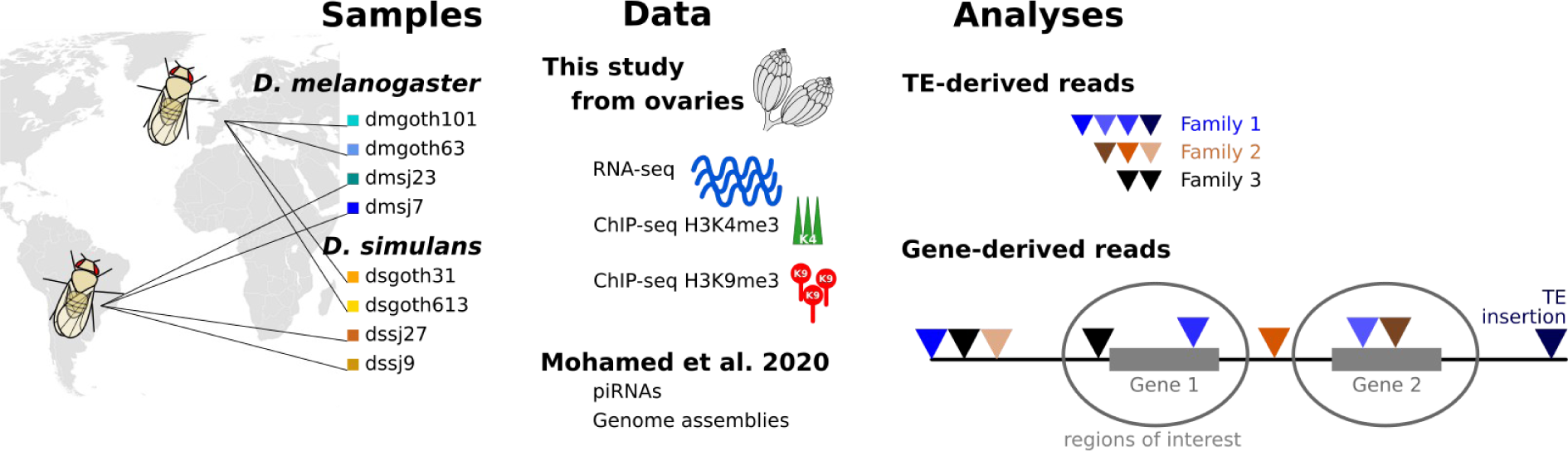
Graphic summary of the study. Eight wild-type strains from *D. melanogaster* and *D. simulans* were included in the study. The present datasets are RNA-seq and ChIP-seq for H3K4me3 and H3K9me3 marks, and were prepared from ovarian samples. They were analyzed in parallel with already published data produced from the same *Drosophila* strains: ovarian small RNA repertoires and genome assemblies based on Oxford Nanopore long read sequencing (Mohamed et al. 2020). For RNA-seq and ChIP-seq, TE-derived reads were analyzed at the TE family level, and gene-derived reads were analyzed in relation to TE insertions inside or near genes (therefore restricted to the TE insertions included within the gray bubbles).

The original approach and subsequent analyses reveal a stronger epigenetic influence of TEs on orthologous genes in *D. simulans* compared to *D. melanogaster*, and are in agreement with the recent work published by Lee’s lab (Huang et al. 2022). At the same time, we uncover a larger contribution of TEs to genome architecture in *D. melanogaster*: in particular, TE insertions contribute more to gene H3K9me3 level variance in *D. melanogaster* compared to *D. simulans*, which is evidenced by a stronger association of TEs around genes with the levels of this chromatin mark in *D. melanogaster*. Overall, this work contributes to the understanding of species-specific influence of TEs within genomes. As a whole, these results participate in the accurate, quantitative understanding of TEs’ impacts on genomes, and highlight the species-specific differences in the interaction between TEs and the host genome. This sheds a new light on the considerable natural variability resulting from TEs, which may be associated with contrasted adaptive and evolutionary potentials, all the more important in a rapidly changing environment (Baduel et al. 2021; Fablet & Vieira 2011; Mérel et al. 2021).

## Results

### TE expression and epigenetic targeting in *Drosophila* ovaries

We first considered TE-derived RNA-seq reads from all samples, which we analyzed at the TE-family level (Figure 1). As performed by other research studies (Chakraborty et al. 2021; Kofler, Nolte, et al. 2015), we removed the non-autonomous *DNAREP1* helentron (also known as *INE-1*) from our analyses because it is a highly abundant element displaying mainly fixed insertions in the *melanogaster* complex of species (Thomas et al. 2014). However, a recent study revealed an expansion of this family in the *Drosophila nasuta* species group (Wei et al. 2022), indicating its activity and potential genomic impacts. We therefore performed a *DNAREP1*-dedicated analysis, apart from the other families. TEs account for 0.6% (dmgoth101) to 1.2% (dmsj23), and 0.5% (dssj9) to 0.7% (dssj27), of read counts corresponding to annotated sequences (genes and TEs) within the ovarian transcriptomes of *D. melanogaster* and *D. simulans* strains, respectively (Figure 2A). However, more gene sequences are annotated in the *D. melanogaster* genome compared to *D. simulans*. Therefore, we also performed the same computation using only 1:1 orthologs, and found similar trends: TEs represent 0.9% (dmgoth101) to 1.8% (dmsj23), and 0.7% (dssj9) to 1.0% (dssj27) of these read counts (Supplemental Fig. S1A). *DNAREP1* accounts for 6% to 13%, and for 5% to 9% of the total number of TE read counts in *D. melanogaster* and *D. simulans*, respectively. This contribution is very weak with regard to the ∼4,000 copies of *DNAREP1* identified by our procedure within each genome. We removed *DNAREP1* and found significant positive correlations between per TE family RNA counts and family genomic coverage (quantified as the total number of bp spanned by each TE family along the genome) (Spearman correlations, rho = 0.33 to 0.37, and 0.39 to 0.44, in *D. melanogaster* and *D. simulans*, respectively; Supplemental Fig. S2A). Regarding TE-derived piRNA production, it was previously described in control conditions in wild-type strains that the amounts of piRNAs were positively correlated with the amounts of RNAs, at the TE family level (Lerat et al. 2017). This remains true in the present dataset: we find significant positive correlations between per TE family RNA counts and piRNA counts (Spearman correlations, rho = 0.39 to 0.48, and 0.48 to 0.56, in *D. melanogaster* and *D. simulans*, respectively; Supplemental Fig. S2B). In both cases, correlations are significantly stronger in *D. simulans*, compared to *D. melanogaster* (Wilcoxon rank tests for *D. melanogaster vs D. simulans* comparisons; correlation coefficients between TE RNA counts and TE sequence abundance: p-value = 0.029; correlation coefficients between TE RNA counts and TE piRNA counts: p-value = 0.029), suggesting a more efficient production of TE-derived piRNAs.

**Fig. 2.**
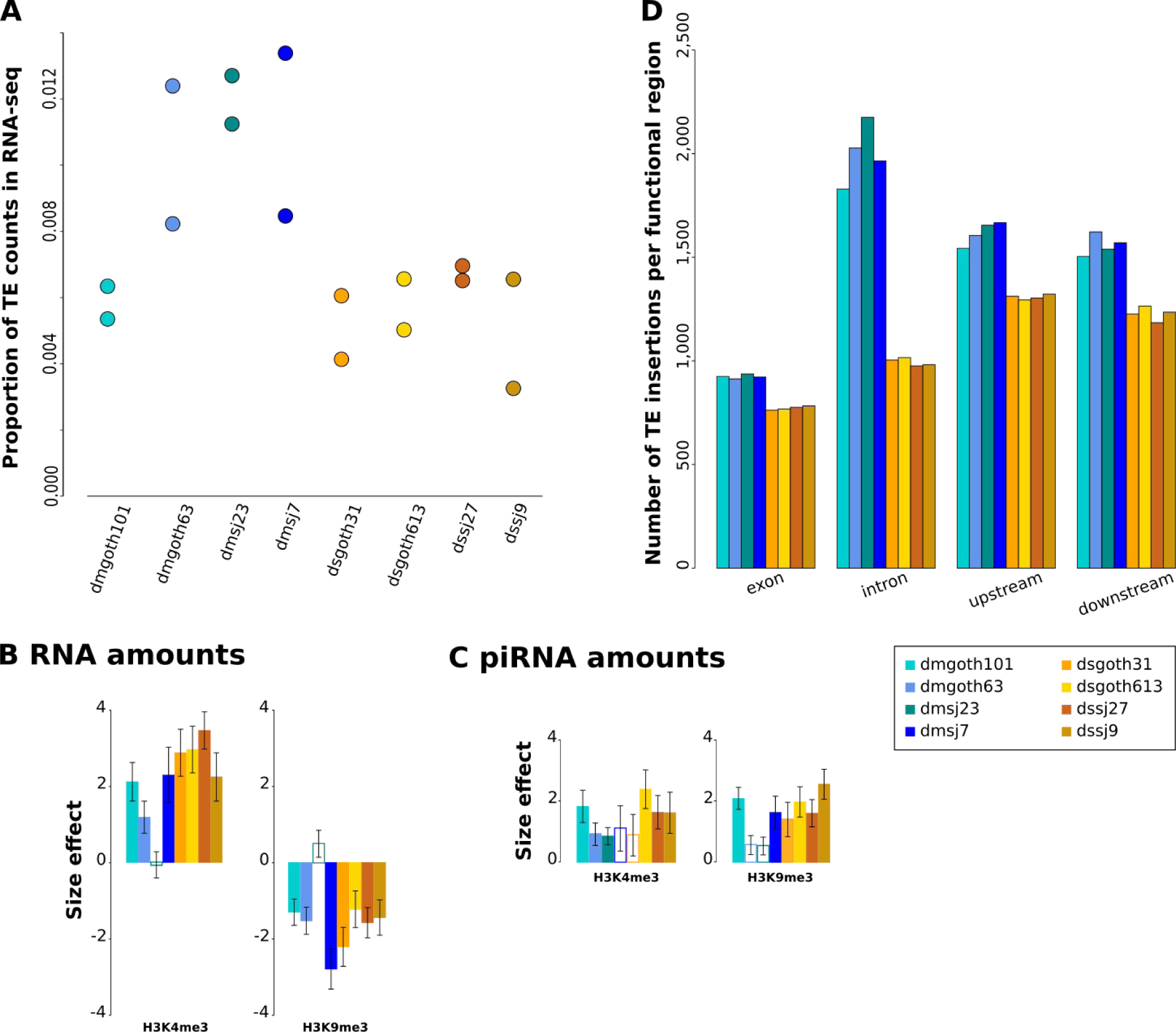
TE insertions and expression. (*A*) Proportions of TE read counts in RNA-seq data relative to read counts corresponding to genes and TEs. For each strain, two biological replicates are shown. (*B*) Contributions of H3K4me3 and H3K9me3 enrichment to TE-derived RNA read counts (according to the model RNA ∼ H3K4me3 + H3K9me3 + input calculated on log10 transformed read count numbers, at the TE family level). Colored bars: p-values < 0.05, empty bars: p-values > 0.05. Error bars are standard errors. (*C*) Contributions of H3K4me3 and H3K9me3 enrichment to TE-derived piRNA read counts (according to the model piRNA ∼ H3K4me3 + H3K9me3 + input calculated on log10 transformed read count numbers, at the TE family level). Colored bars: p-values < 0.05, empty bars: p-values > 0.05. Error bars are standard errors. (*D*) Number of TE insertions per functional region per strain. Upstream and downstream regions are 5 kb sequences directly flanking transcription units 5’ and 3’, respectively.

We assessed the contribution of histone mark enrichment to TE RNA amounts considering the following linear model on log-transformed normalized read counts: RNA ∼ H3K4me3 + H3K9me3 + input. These models led to adjusted r^2^ as high as 0.48 to 0.64 depending on the strains in *D. melanogaster* and 0.45 to 0.60 in *D. simulans*, suggesting that these models capture significant portions of TE RNA amount variation. We find that TE RNA amounts are positively correlated with H3K4me3 and negatively correlated with H3K9me3 amounts (Figure 2B), as expected considering that H3K4me3 is an activating mark while H3K9me3 is a silencing one. Input amount contributions to r^2^ are very low (< 0.05, Supplemental Table S3). We used a similar approach to analyze piRNA amounts, and considered the following linear model on log-transformed read counts: piRNA ∼ H3K4me3 + H3K9me3 + input. We obtained even higher adjusted r^2^ values, from 0.70 to 0.75, and 0.64 to 0.68, depending on the strains in *D. melanogaster* and *D. simulans*, respectively. We find that TE-derived piRNA amounts are positively correlated both with permissive H3K4me3 and repressive H3K9me3 levels (Figure 2C). The tighter correlations may be due to the strong dependency of piRNA production mechanisms on chromatin marks and H3K9me3 in particular, while RNA transcription also involves other factors, such as transcription factors, whose binding sites vary a lot across TE sequences.

### TE insertions within or nearby genes

In the following sections, we focus on gene-derived reads from all samples, which we analyzed with regard to the presence of TE insertions within or nearby genes (Figure 1). Based on gene annotations, we distinguished the different functional regions of genes: exons, introns, upstream, or downstream sequences (5 kb flanking regions). Exons are both UnTranslated Regions (UTRs) and Coding Sequences (CDSs). Sequences that may both behave as exons or introns depending on alternative splicing are included in “exons”. In this first step, we considered a set of 17,417 annotated genes for *D. melanogaster*, and 15,251 for *D. simulans* (see Material and Methods). We quantified the number of TE insertions within genes (Figure 2D), and found that they account for ∼25% and ∼16% of the total number of TE insertions per genome in *D. melanogaster* and *D. simulans*, respectively. The lower proportion observed in *D. simulans* for TE insertions retained within genes suggests a stronger selection against TE insertions in this species compared to *D. melanogaster*, assuming that other genomic characteristics are similar. This difference holds true when considering only the 12,470 1:1 orthologs between *D. melanogaster* and *D. simulans* (Supplemental Fig. 1B). Among the copies of *DNAREP1* that we identified along the genomes, our analysis revealed that 1,343 to 1,374 insertions from this family are found within genes in *D. melanogaster*, and 1,075 to 1,089 insertions in *D. simulans*.

### TE insertions are associated with variability in expression and histone enrichment between ortholog genes

We used our experimental dataset to infer the contribution of TE insertions at the inter-genomic level, *i.e.* we compared expression levels of the same genes across genomes. We focused on the subset of genes that we found expressed in the ovaries (see Material and Methods), *i.e.* 7,883 to 8,135 genes depending on the strains of *D. melanogaster*, and 7,653 to 8,121 genes in *D. simulans*. We first considered *D. melanogaster* and *D. simulans* separately. For each gene that displays variation in TE insertion numbers across strains, we computed the mean difference of gene expression (TPM, scaled by gene average) between the strain that had the highest TE insertion numbers and the strain that had the lowest. When several strains had the same numbers of TE insertions, we computed their average gene expression level. We performed the same approach on histone enrichment. Our assumption was that a general effect of TE insertions would shift the distribution of the mean difference away from 0. This is not what we observed for RNA levels nor for H3K4me3 enrichment (one-sample t tests, all p-values > 0.05) (Figure 3, Supplemental Fig S4). However, we find an increase in H3K9me3 enrichment associated with high TE insertion numbers, but only in *D. simulans* and for TE insertions within introns and upstream of genes (one-sample t test; within introns: mean difference = 0.003, p-value = 0.0005; upstream: mean difference = 0.003, p-value = 0.0019). These results are congruent with recent studies, which observed a clear association between TE insertions and heterochromatin but no predominant negative impact on the expression of neighboring genes (Huang et al. 2022; Wei et al. 2022).

**Fig. 3.**
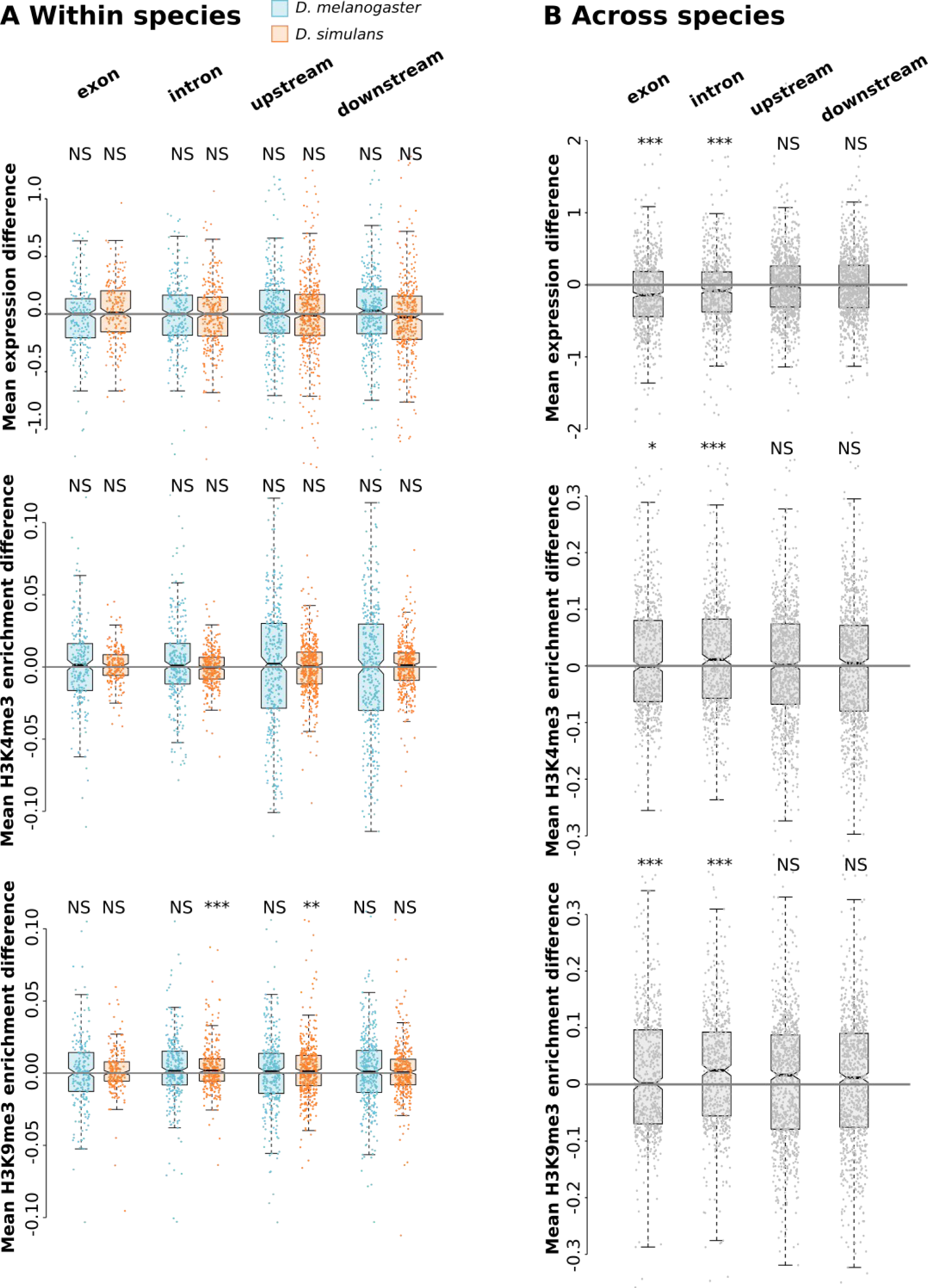
Variability in gene expression and histone enrichment according to TE insertion numbers across strains. (*A*) Mean expression difference (in TPM, scaled by gene average) between strains with the highest and the lowest TE insertion numbers for each region of each gene; mean histone enrichment difference (log-transformed, scaled by gene average) between strains with the highest and the lowest TE insertion numbers. Analyses are performed separately for both species (blue: *D. melanogaster*, orange: *D. simulans*), only considering genes that show different TE insertion numbers across strains. Significance levels correspond to t tests comparing observed mean to 0. (*B*) Same analyses across all eight strains considering 1:1 ortholog genes. Significance levels correspond to t tests comparing observed mean to 0: p-value 0 *** 0.001 ** 0.01 * 0.05.

We also took the opportunity to consider 1:1 ortholog genes (6,417 genes) so as to include all eight strains (*D. melanogaster* and *D. simulans*) in the same analysis (Figure 3B). Computation strategies were the same as above and revealed significant decreases in RNA levels for strains with the highest TE insertion numbers in exons (mean difference = -0.129, p-value = 1e-10) and introns (mean difference = -0.077, p-value = 9e-5). We also found significant increase in H3K4me3 levels as well as H3K9me3 levels for strains with the highest TE insertion numbers in exons and introns (H3K4me3, TEs within exons: mean difference = 0.012, p-value = 0.0201; within introns: mean difference = 0.019, p-value = 1e-5; H3K9me3, TEs within exons: mean difference = 0.037, p-value = 0.0092; within introns: mean difference = 0.028, p-value = 2e-5). However, such an analysis including all strains from both species at once has to be considered with caution because gene sequences differ across species (GC content, length, etc.), which may interfere with mapping and read counting, and was not accounted for in this work. In addition, the comparisons may be confounded by genome-wide differences in TE density –globally higher in *D. melanogaster*– or H3K9me3 levels –globally higher at TE insertions in *D. simulans*.

### TE insertions are associated with RNA level variability across genes within genomes

One of the novelties of the present work is to quantify the contribution of TE insertions to the variance in gene expression levels within distinct genomes. Again, we focused on the subset of genes that we found expressed in the ovaries. We quantified TE insertion contribution to gene RNA levels using the following linear models built on log-transformed TPM (Transcript Per Million): TPM ∼ exon + intron + upstream + downstream, where these variables correspond to the number of TE insertions within exons, introns, 5 kb upstream, and 5 kb downstream regions, respectively. We find that TE insertions contribute significantly, albeit weakly, to gene expression variance (Figure 4A): 1.6% to 1.9% of total variance in *D. melanogaster*; 1.2% to 1.9% in *D. simulans*. These values may look low at first sight; however, gene expression levels are known to be primarily regulated by many other factors, such as transcription factor binding, sequence composition and polymorphism, etc. This reveals that our approach is powerful enough to capture low levels of variation and that TEs are significant actors of this variability. Although total contribution to gene expression variance does not differ between species (Wilcoxon rank test, p-value = 0.685), we found significant differences when considering specific gene regions. For instance, the contribution of TE insertions within introns was higher in *D. simulans* compared to *D. melanogaster* (mean values: 0.03% *vs* 0.14%; Wilcoxon rank test, p-value = 0.029), while the contribution of TE insertions downstream of genes was higher in *D. melanogaster* compared to *D. simulans* (mean values: 0.06% *vs* 0.21%; Wilcoxon rank test, p-value = 0.029).

**Fig. 4.**
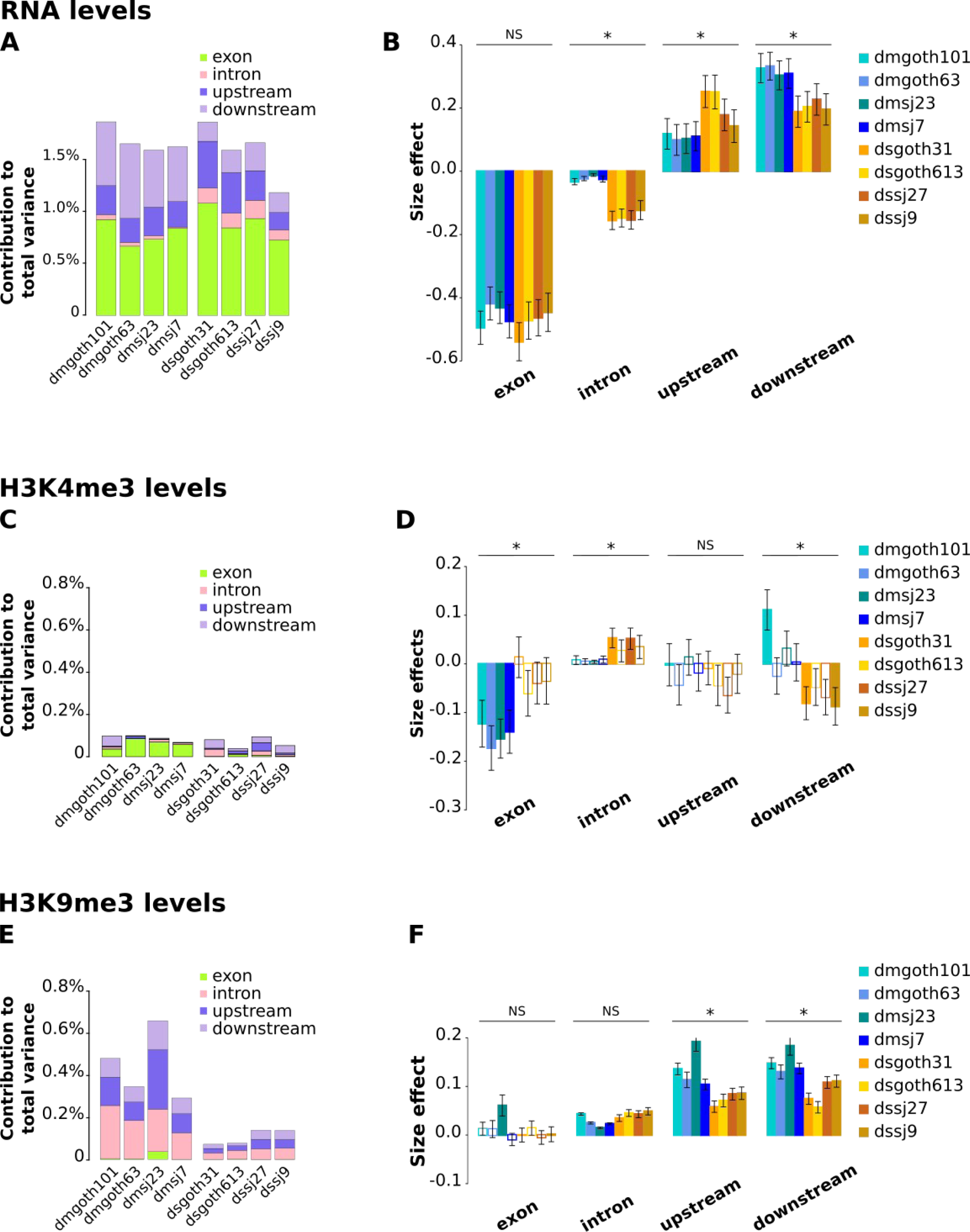
TE insertions are associated with RNA levels and histone enrichment variability across genes within genomes. (*A*) Contribution of TE insertion numbers to gene expression total variance estimated using the linear model gene TPM (log) ∼ exon + intron + upstream + downstream, and (*B*) corresponding size effects. (*C*) Contribution of TE insertion numbers to gene H3K4me3 total variance estimated using the linear model gene H3K4me3 level (log) ∼ exon + intron + upstream + downstream, and (*D*) corresponding size effects. (*E*) Contribution of TE insertion numbers to gene H3K9me3 total variance estimated using the linear model gene H3K9me3 level (log) ∼ exon + intron + upstream + downstream, and (*F*) corresponding size effects. Significance indications above graphs in *(B, D, E)* are *D. melanogaster vs D. simulans* comparisons using Wilcoxon rank tests. Colored bars: p-values < 0.05, empty bars: p-values > 0.05. Error bars are standard errors.

When we computed the corresponding size effects, we observed significant, negative associations between gene expression levels and TE insertions within exons and introns, and significant, positive associations for TE insertions around genes (Figure 4B). The association with gene expression was stronger for *D. melanogaster* compared to *D. simulans* for downstream TE insertions (Fold-change = 1.6; Wilcoxon rank test, p-value = 0.029), and it was stronger in *D. simulans* compared to *D. melanogaster* for TE insertions within introns (Fold-change = 6.2; Wilcoxon rank test, p-value = 0.029) and upstream TE insertions (Fold-change = 1.9; Wilcoxon rank test, p-value = 0.029).

Nevertheless, one could argue that the species-specific differences that we observe here are due to gene sets not being exactly the same across species. In order to correct for this bias, we focused on the subset of 6,417 genes that have 1:1 ortholog in the other species and that are expressed in ovaries. The results were very similar regarding size effects, reinforcing our conclusions (Supplemental Fig. S5). However, we noticed that TE contribution to gene expression variance was increased in this subset of genes: 3.2% and 2.9% on average in *D. melanogaster* and *D. simulans*, respectively (Supplemental Fig. S5).

Collectively, our data show a weak but significant contribution of TEs to the variance in gene expression within genomes, which varies across species and is due to negative correlations between gene RNA levels and TE numbers in exons and introns, and positive correlations with TE numbers upstream and downstream of genes.

### TE insertions are associated with histone enrichment variability across genes within genomes

We used a similar approach to analyze H3K4me3 and H3K9me3 enrichment (*i.e.* we aligned ChIP-seq reads against whole gene sequences and computed corresponding read counts), and used the models H3K4me3 or H3K4me3 ∼ input + exon + intron + upstream + downstream. We found that TE insertions contributed significantly (except in dsgoth613), albeit very weakly, to gene H3K4me3 levels variance (0.07% to 0.10% total variance in *D. melanogaster*; 0.04% to 0.09% in *D. simulans*; Wilcoxon rank test for *D. melanogaster vs D. simulans* comparison, p-value = 0.200) (Figure 4C). When computing size effects, the only significant and consistent result is a negative association of TE insertions within exons with gene H3K4me3 levels, in *D. melanogaster* only (Figure 4D).

The contribution of TE insertions to total variance is higher for H3K9me3 levels: 0.29% to 0.65% in *D. melanogaster*, and 0.07% to 0.14% in *D. simulans* (Figure 4E; Wilcoxon rank test for *D. melanogaster vs D. simulans* comparison, p-value = 0.029). The largest contribution comes from TE insertions around genes and within introns, while TE insertions within exons virtually do not contribute to H3K9me3 variance. The computation of size effects reveals a consistent, positive association of TE insertions within introns, upstream and downstream genes with H3K9me3 levels, in both species. These results are in agreement with TEs being the preferential targets for H3K9me3 deposition, which then spreads to neighboring regions (Le Thomas et al. 2013; Rebollo et al. 2011). Alternatively, we cannot exclude that they may also lie in particular chromatin environments where there is retention bias (Sultana et al. 2017), and that the associations detected here are due to these particular chromatin features. The effects are stronger in *D. melanogaster* compared to *D. simulans* for TE insertions around genes (Figure 4F; Upstream: fold-change = 1.8, Wilcoxon rank test, p-value = 0.029; Downstream: fold-change = 1.7, Wilcoxon rank test, p-value = 0.029).

When considering only the set of 1:1 orthologous genes, patterns are highly similar for size effects, except that the association between TE insertions within introns and H3K9me3 levels is now significantly stronger in *D. melanogaster* compared to *D. simulans.* In addition, the contribution to H3K4me3 total variance is higher for this subset of genes compared to the total set, although it remains very low, up to 0.73% in *D. melanogaster* and 0.37% in *D. simulans*. (Supplemental Fig. S5).

While the observation of concomitant negative correlations with RNA levels and positive correlations with H3K9me3 for TE insertions within introns is in agreement with a negative impact of a heterochromatic mark on gene expression, the results for TE insertions around genes appears a little bit at odds. Indeed, TE insertions upstream and downstream of genes are at the same time positively correlated with RNA levels and H3K9me3 enrichment. One hypothesis for these TE insertions could be that their positive association with RNA levels is due to the multiple transcription factor binding sites that they bring —some transcription factors such as CTCF are known to be insensitive to chromatin (Isbel et al. 2022)—, and this ends up counteracting the negative impact of H3K9me3 targeting.

### Patterns are globally conserved across TE classes and ages

We next analyzed TE insertions according to TE class, *i.e.* LTR (Long Terminal Repeat) elements, LINEs (Long Interspersed Nuclear Elements), DNA transposons, and *DNAREP1*. We used the same linear models on the same sets of genes, but considering only TE insertions belonging to each particular class. TE insertion numbers vary across classes (Supplemental Fig. S6), which leads to differences in statistical power (the higher power associated with the higher number of TE insertions). Despite this, the computation of size effects on gene RNA levels, H3K4me3, and H3K9me3 levels revealed highly consistent patterns across TE classes (Supplemental Fig. S6). *DNAREP1* patterns are similar to other DNA transposons. The major difference with global patterns (Figure 4) is a trend for a positive association of DNA transposons and *DNAREP1* insertions in exons with gene expression in *D. melanogaster* only. Differences between transposons (DNA transposons and *DNAREP1*) and retrotransposons (LTR elements and LINEs) might be related to different waves of transposition: Kofler *et al*. described that LTR insertions are mostly of recent origin in both species, while DNA and non-LTR insertions are older, and that DNA transposons showed higher activity levels in *D. simulans* (Kofler, Nolte, et al. 2015). The positive association between TE insertions in exons and gene expression would be characteristics of the families with the most ancient transposition activity, and potentially domestication events.

Irrespective of TE classes, it has already been described that TEs’ impacts on genes differ across young (*i.e.* polymorphic) and old (*i.e.* fixed) TE copies; this is due to the pool of old TE insertions having been purged from deleterious insertions by natural selection (Hollister & Gaut 2009). Indeed, Uzunovic *et al*. (Uzunović et al. 2019) showed in the plant *Capsella* that young TE insertions had a negative effect on gene expression while old insertions were more likely to increase gene expression. In this view, we distinguished insertions that are unique to one genome and absent in the three other strains of the same species (“private”) —and therefore correspond to the most recent insertions—, and those that are shared by all four strains of the species (“common”) —thus the oldest ones. Many of the TE insertions that are considered here (61% in *D. simulans* to 64% in *D. melanogaster*) fall in the “common” category. This may seem at odds regarding previous knowledge and the work of Kofler *et al*. in particular, who found that >80% TE insertions had low frequency in pool seq data (Kofler, Nolte, et al. 2015). However, the majority of these insertions are intergenic while we only focus on TEs within or around genes in the present study, which explains the differences in proportions between the two studies. The difference in subset sizes between “common” and “private” categories also leads to a reduced statistical power for the set of private insertions. Despite this difference, the observed patterns are rather consistent between both sets of TEs, and very similar to the global patterns including all TEs regardless of insertion polymorphism (Figure 4, Supplemental Fig. S7). In the “common” pool, we do not observe the positive association between TE insertions in exons and gene expression reported by (Uzunović et al. 2019), maybe because the majority of these insertions are not old enough, or at least not as old as the above-described DNA transposon pool in *D. melanogaster*. Since our approach is gene-centered (Figure 1), it is very likely that our complete set of TE insertions is already biased: when deleterious, insertions within or near genes have such a negative impact that we are not able to catch them from natural samples. Therefore, our complete set of TE insertions may already correspond to copies that have passed the filter of natural selection, and thus does not show critical differences between “common” and “private” patterns. However, some species-specific difference appears in the private set of insertions within introns: they display stronger negative association with gene expression levels in *D. simulans*, and stronger positive association with H3K9me3 levels in *D. melanogaster*. We speculate that this reveals species-specific differences in the efficiency of TE control at the first stages of TE invasion.

### Gene-derived small RNAs and epigenetic effects

It has been demonstrated that TEs are sources of piRNA biogenesis in the ovary through the action of Rhino that promotes non-canonical transcription (Mohn et al. 2014). We took advantage of our extensive dataset composed of RNA-seq, ChIP-seq and small RNA-seq produced from the ovaries of the exact same strains to test for the impact of piRNA cluster activity on neighbouring genes. In addition, siRNAs were previously shown to be produced from piRNA clusters and participated in TE silencing in ovaries (Shpiz et al. 2014). Therefore, we searched for gene-derived piRNAs and siRNAs, which could result from the spreading of small RNA production machinery from TE insertions. We filtered small RNAs based on read length, which does not allow us to distinguish siRNAs from miRNAs in the pool of 21 nt reads. We will therefore refer to them as “21 nt RNAs”. In agreement with this scenario, we found a significant positive correlation between gene-derived piRNAs and gene-derived 21 nt RNAs (Spearman correlation coefficients; *D. melanogaster*: 0.517 to 0.536; *D. simulans*: 0.526 to 0.661; all p-values < 1e-10). In addition, we found that gene-derived piRNA production was significantly positively correlated with gene H3K9me3 levels (Supplemental Fig. S8) (Spearman correlation coefficients; *D. melanogaster*: 0.561 to 0.586; *D. simulans*: 0.475 to 0.525; all p-values < 1e-10). These results are compatible with a scenario of piRNA cluster transcription spreading to nearby gene sequences. Remarkably, correlations were stronger for *D. melanogaster* compared to *D. simulans* (Wilcoxon rank test, p-value = 0.029). Gene-derived 21 nt RNA production was also significantly positively correlated with gene H3K9me3 (Spearman correlation coefficients; *D. melanogaster*: 0.470 to 0.517; *D. simulans*: 0.437 to 0.504; all p-values < 1e-10) but the strength of the correlation was not significantly different between species.

In addition, in order to test whether TEs could be driving this correlation between gene-derived piRNAs and H3K9me3 levels, we focused on expressed genes whose polymorphic TE insertions were only “private”. These TE insertions are assumed to be the most recent and therefore the ones with the strongest epigenetic spreading. We found that piRNA production from these genes were more frequently higher than the third quartile than expected (Supplemental Table S9). These results demonstrate that the control of TE sequences by the piRNA pathway impacts neighboring genes through the production of gene-derived small RNAs and the increased deposition of H3K9me3 marks.

## Discussion

The common-held view is that, as parasites that are fought against by genomes, TEs have a general negative impact on gene expression (Cridland et al. 2015; Lee & Karpen 2017; Lee 2015). Our present findings are in agreement with this idea. However, the originality of this research work is to provide an unprecedented quantitative view, which allows to precisely decipher TE impacts, integrating data gathered from wild-type strains of two closely related *Drosophila* species. This study combines genomic, transcriptomic, and epigenetic high-throughput sequence data, all produced from ovaries, where TEs are tightly controlled by epigenetic mechanisms through the piRNA pathway (Malone & Hannon 2009; Senti & Brennecke 2010) and therefore where we are to expect the strongest impacts of TEs on genes.

### Expression and epigenetic marks of TE sequences

Our results uncover a lower contribution of TEs to the *D. simulans* transcriptome as compared to *D. melanogaster* (0.6% *vs* 1.1% on average, Figure 2A). This is in agreement with the previously described lowest contribution of TEs in the genomes of *D. simulans* in terms of sequence abundance and copy numbers (Mohamed et al. 2020; Vieira et al. 1999). However, these figures are not proportional to TE abundances in the genomes of both species (12.2% in *D. simulans vs* 19.3% in *D. melanogaster* (Mérel et al. 2020)) and indicate a stronger inhibition of TE expression in *D. simulans* compared to *D. melanogaster*. In both species, we found that H3K9me3 marks on TE sequences are associated with a decrease in TE-derived RNA amounts, and the opposite for H3K4me3 marks. On the contrary, we observed that both histone marks are positively correlated with TE-derived piRNA amounts, which is congruent with the piRNA-targeted deposition of H3K9me3 marks at transcriptionally active TE copies (Czech et al. 2018; Sienski et al. 2012). However, one should note that these results reflect average behaviors at the TE family level, and TE copies may differ from one another within TE families.

What emerges from the different analyses that we performed is a remarkable variability across TEs, as illustrated by the width of dot distributions in Figure 3 for instance. This highlights the huge variability across TE sequences on many aspects: class, family, length, insertion site preference, chromosome distribution, activity, transposition rate, etc. For instance, in their pool-seq analysis of *D. melanogaster* and *D. simulans*, Kofler *et al*. found that half of the TE families showed evidence of variation of activity through time and were not the same depending on the species (Kofler, Nolte, et al. 2015). It is congruent with the conclusions of Wei *et al*., working on the *Drosophila nasuta* complex of species, who emphasize that TE insertions can have multiple effects on gene expression, from no effect to silencing or over-expression (Wei et al. 2022). This also echoes the work of Malone *et al*. and Sienski *et al*., who described different groups of TEs depending on their sensitivity to different piRNA pathways and thus different effects on neighboring genes (Malone et al. 2009; Sienski et al. 2012). In addition, it has already been suggested and demonstrated that TEs’ influence on gene expression is only manifested in case of stress (Naito et al. 2009), which adds another layer of variability and difficulty to disentangle biological impacts.

### Intra- and inter-genomic analyses tell distinct, although complementary stories

In the intra-genomic analysis, we gather all expressed genes from a given genome, which we compare for their TE insertions, expression level, chromatin marks, and piRNA production. These are therefore heterogeneous sets of genes, which work coordinately in living cells. In the inter-genomic analysis, we compare the same ortholog genes in different genomes. We assume that these genes differ mainly based on their TE insertions.

When TE insertions are associated with differences in gene expression or chromatin state, it is very difficult to tell apart whether these TE insertions are causative or not. Nevertheless, the inter-genomic analysis is a way to demonstrate causality because it compares versions of the same genes but displaying different numbers of TE insertions —however with the limitation of neglecting nucleotide polymorphism. This approach has already successfully been followed by others and led to the conclusion of the causative role of the TE insertions (Lee & Karpen 2017; Rebollo et al. 2011). On the contrary, in the intra-genomic study, we draw general patterns from the analysis of the complete set of genes at once, which differ from TE insertion numbers but also from many other aspects (sequence, length, expression level, tissue-specificity, local recombination rate, etc.). The intra-genomic analysis allows to identify associations between TE insertions, gene expression and chromatin environment, and therefore brings us to draw species-specific gene landscapes.

Here, the inter-genomic analysis on the complete dataset (orthologous genes from both species, Figure 3B) reveals that TE insertions within, but not around genes, have a negative impact on gene RNA levels, and a positive impact on both histone marks, H3K4me3 and H3K9me3. This H3K4me3 result may be related to TEs donating promoters or *cis* regulatory sequences, as was already described on several instances (Moschetti et al. 2020; Sundaram et al. 2014; Villanueva-Cañas et al. 2019) or disrupting inhibitory sequences. The impact on H3K9me3, however, appears to be stronger since the net result is negative on gene RNA levels. This result corresponds to TEs being a preferential target for H3K9me3 deposition (Le Thomas et al. 2013), which then spreads to neighboring sequences. However, this pattern could also be due to some disruptive effects caused by TE insertions in the gene body, which would then reduce the selective pressures to maintain expression levels as high as their initial levels.

In addition, the inter-genomic analysis reveals stronger epigenetic impacts of TE insertions in *D. simulans* compared to *D. melanogaster* (Figure 3A). These results support the previous findings from Lee & Karpen, which found higher enrichment and spread of H3K9me2 from TE insertions in *D. simulans* compared to *D. melanogaster (Lee & Karpen 2017)*. These results were recently confirmed in a larger set of species (Huang et al. 2022). They proposed that this leads to stronger selection against TE insertions close to genes in *D. simulans* compared to *D. melanogaster*, which explains the lower total number of TE insertions and the lower proportion of TE insertions within or nearby genes in *D. simulans*. However, even if we were able to detect mean effects of TE insertions, our results also reveal a large variety of impacts of individual TE insertions —as illustrated by the width of dot distributions in Figure 3 for instance—, either positive or negative, which suggests that TE effects may not be as pervasive as previously claimed (Lee & Karpen 2017).

On the other hand, the intra-genomic analysis confirms the already described trend of TE insertions within genes to be associated with a reduction in gene RNA levels. However, our results also reveal that TE insertions around genes are associated with increased gene expression on average. Overall, TE insertions are virtually not associated with particular H3K4me3 patterns, except for TE insertions in exons in *D. melanogaster*, which are associated with a decrease in H3K4me3. As previously known and confirmed by the inter-genomic analysis, TE insertions are associated with increased levels of H3K9me3. The novelty brought by the intra-genomic analysis is that the association is particularly strong for TE insertions around genes and not within genes, particularly in *D. melanogaster* compared to *D. simulans*. *D. melanogaster* TEs contribute more to gene H3K9me3 level variance compared to *D. simulans*. This suggests that there is stronger structuring or stratification of genes according to TE insertion numbers and histone marks in this species compared to *D. simulans*. TE insertions are more frequently found with higher H3K9me3 (and even H3K4me3 to a lesser extent) enrichment in *D. melanogaster*.

Interpretations from inter- and intra-genomic analyses seem contradictory at first sight. However, they may illustrate the two facets of RNA interference, *i.e.* defense *vs* regulation (Torri et al. 2022). We may speculate that in *D. simulans*, the defense facet appears prominent while the regulation prevails in *D. melanogaster*. Such differences in closely related species are not unexpected in the piRNA pathway, which is known to be evolving at a particularly elevated rate (Fablet et al. 2014; Obbard et al. 2009). Again, we may speculate that this is related —whether as a cause or a consequence cannot be told— to the different tempo of TE activity and genome colonization between both species.

In the intra-genomic analysis, many parameters other than the numbers of TE insertions differ across the genes (the family and length of the TEs, gene sequence composition, presence of transcription factor binding sites, etc. (Hill et al. 2021; Wittkopp & Kalay 2011)) and yet we were able to capture statistical signal from the numbers of TE insertions. This suggests a widespread influence of TEs on gene expression. The underlying mechanisms may be chromatin mark spreading, but not only. TEs may also disrupt functional elements, especially for those inside genes, or add transcription factor binding sites (Horváth et al. 2017; Rebollo, Romanish, et al. 2012; Ullastres et al. 2021). Moreover, we have to note that TE insertions may accumulate in specific chromatin environments due to insertional preference or different levels of selection in these environments (Sultana et al. 2017).

### TEs’ influence on genomes is contrasted between *D. melanogaster* and D. simulans

The intra- and inter-genomic analyses performed here both reveal species-specific differences, however not at the same scale. The inter-genomic analysis reveals a stronger epigenetic inhibition of TE sequences in *D. simulans* compared to *D. melanogaster*, indicative of a stronger counter-selection of TE insertions. In parallel, the intra-genomic analysis uncovers stronger associations between epigenetic landscape and TE insertions in *D. melanogaster,* and a positive association between gene expression and TE insertions located in the flanking regions (Figure 4). It means that genes that have many TEs in *D. melanogaster* on average have higher H3K9me3 levels than genes that have many TEs in *D. simulans*. This may be due to differences in TE insertion landscapes or to differential retention in particular chromatin regions. This analysis therefore reveals how TE sequences may participate in the structure of the genome and how this differs between species. This reflects more long-term and intimate interactions between the host genome and its TEs.

The species-specific differences that we observe for TE influence on genes may be due to variability in the efficiency of epigenetic machinery, as suggested by (Lee & Karpen 2017; Rebollo, Horard, et al. 2012). Alternatively, it may also reveal different tempo of TE dynamics between these species. A recent peak of activity of TEs can be seen in *D. melanogaster*, which is much smaller in *D. simulans (M*érel et al. 2020), indicating that the colonization of the *D. simulans* genome by TEs started more recently (as suggested by our previous results (Mohamed et al. 2020) and others (Kofler, Hill, et al. 2015)). Such ongoing colonization would also lead to the selection of more efficient TE control mechanisms.

These contrasted impacts of TE insertions on genes through epigenetic marks across the species provide an additional demonstration of the considerable natural variability due to TEs. We predict that this leads to contrasted adaptive and evolutionary potentials, all the more determining in a rapidly changing environment (Baduel et al. 2021; Fablet & Vieira 2011; Mérel et al. 2021).

## Material and Methods

### Drosophila strains

The strains under study in the present work were previously described in Mohamed *et al*. (Mohamed et al. 2020). The eight samples of *D. melanogaster* and *D. simulans* wild-type strains were collected using fruit baits in France (Gotheron, 44°56’0”N 04°53’30”E - “goth” strains) and Brazil (Saõ Jose do Rio Preto 20°41’04.3”S 49°21’26.1”W – “sj” strains) in June 2014. Two isofemale lines per species and geographical origin were established directly from gravid females from the field (French *D. melanogaster*: dmgoth63, dmgoth101; Brazilian *D. melanogaster*: dmsj23, dmsj7; French *D. simulans*: dsgoth613, dsgoth31; Brazilian *D. simulans*: dssj27, dssj9). Brothers and sisters were then mated for 30 generations to obtain inbred strains with very low intra-line genetic variability. Strains were kept at 24°C in standard laboratory conditions on cornmeal–sugar–yeast–agar medium.

### Genome annotation

Genome assemblies were produced in (Mohamed et al. 2020) and have been deposited in the European Nucleotide Archive (ENA) at EMBL-EBI under accession number PRJEB50024 (https://www.ebi.ac.uk/ena/browser/view/PRJEB50024). Throughout the present analysis, we kept scaffolds corresponding to complete chromosomes 2L, 2R, 3L, 3R, 4, and X.

TE annotation: We used RepeatMasker 4.1.0 (http://repeatmasker.org/) -species Drosophila in order to identify TE sequences in the assemblies, followed by OneCodeToFindThemAll (Bailly-Bechet et al. 2014) with default parameters, in order to parse RepeatMasker results. We include all TE sequences in the subsequent analyses, whether they are full length or truncated.

Gene annotation: We retrieved gtf files from FlyBase : ftp.flybase.net/genomes/Drosophila_melanogaster/dmel_r6,46_FB2022_03/gft/dmel-all-r6.46.gtf.gz and ftp.flybase.net/genomes/Drosophila_simulans/dsim_r2,02_FB2017_04/gtf/dsim-all-r2,02.gtf.gz. The corresponding fasta files were also downloaded from FlyBase: ftp.flybase.net/genomes/Drosophila_melanogaster/dmel_r6,46_FB2022_03/fasta/dmel-all-chromosome-r6.46.fasta.gz and ftp.flybase.net/genomes/Drosophila_simulans/dsim_r2,02_FB2017_04/fasta/dsim-all-chromosome-r2,02.fasta.gz. We used Liftoff (Shumate & Salzberg 2020) to lift over gene annotations from the references to our genome assemblies. We used -flank 0.2 and only kept the “gene” and “exon” terms. Then, we used the GenomicRanges R package (version 1.38.0) (Lawrence et al. 2013) and the subsetByOverlaps function to cross gene and TE annotations.

1:1 orthologs: We retrieved ortholog information from FlyBase (ftp://ftp.flybase.net/releases/current/precomputed_files/orthologs/dmel_orthologs_in_drosophila_species_fb_2022_01.tsv.gz) and kept only those genes for which there was a 1 to 1 correspondence between *D. melanogaster* and *D. simulans*. TE genomic sequence abundance (bp) was computed using OneCodeToFindThemAll (Bailly-Bechet et al. 2014).

In order to determine which TE insertions were common (shared) to the four strains of a species or unique (private) to one strain, we used the following procedure. We used the GenomicRanges R package and the subsetByOverlaps function to build correspondence between gene and TE annotations of each genome assembly. For each gene and each functional region of each genome assembly for a given species, we extracted the family names of the TE insertions. We define as “common” insertions that are found in the same functional region of the same gene in all the other genome assemblies of the same species. We define as “private” to one strain the insertions that are not found in the same functional region of the same gene in any of the other genome assemblies (in particular, identical TE families are excluded).

### RNA-seq preparation

RNA was extracted from ovaries of 30 three to five day-old females. Two replicates per strain were produced. RNA extraction was carried out using RNeasy Plus (Qiagen) kit following manufacturer’s instructions. After DNAse treatment (Ambion), quality control was performed using an Agilent Bioanalyzer. Libraries were constructed from mRNA using the Illumina TruSeq RNA Sample Prep Kit following manufacturer’s recommendations. Libraries were sequenced on Illumina HiSeq 3000 with paired-end 150 nt reads.

### RNA-seq analysis

TE read counts were computed at the family level using the TEcount module of TEtools (Lerat et al. 2017) and the list of TE sequences available at ftp://pbil.univ-lyon1.fr/pub/datasets/Roy2019/.

Genome sequences from *D. melanogaster* and *D. simulans* were downloaded from FlyBase (dmel-all-chromosome-r6.16.fasta and dsim-all-chromosome-r2.02.fasta) and then masked using RepeatMasker (http://repeatmasker.org/). For each species, we then built a multifasta file of gene sequences using bedtools getfasta (Quinlan & Hall 2010) with gff files available from FlyBase (dmel-all-r6.16.gff and dsim-all-r2.02.gff).

Raw reads were processed using Trimmomatic 0.39 (Bolger et al. 2014) ILLUMINACLIP:TruSeq3-PE.fa:2:30:10 LEADING:3 TRAILING:3 SLIDINGWINDOW:4:20 MINLEN:36, then mapped to genes using HiSat2 (Kim et al. 2019). Alignment files were converted to BAM and sorted using SAMtools (Li et al. 2009), and TPM and effective counts were then computed using eXpress (Roberts et al. 2011).

Quantification of the associations between TE insertions and gene transcript levels: considering only genes expressed in ovaries, we computed mean TPM across replicates and used the following linear models after log transformation: TPM ∼ exon + intron + upstream + downstream, where “exon”, “intron” “upstream”, and “downstream” are the numbers of TE insertions in exons, introns, 5 kb upstream sequences, and 5 kb downstream sequences, respectively. Size effects for each of these factors were then recorded as the coefficients for the explanatory variables. To compute the contribution to total variance, we divided the Sum Square of the corresponding variables by the Total Sum Square, provided by the ANOVA of the linear model.

### ChIP-seq preparation

Chromatin immunoprecipitation was performed using 50 ovary pairs dissected from three to five day old females. Ovaries were re-suspended in A1 buffer containing 60mM KCl, 15mM NaCl, 15mM Hepes, 0,5% Triton and 10mM Sodium butyrate. Formaldehyde (Sigma) was added to a final concentration of 1.8% for secondary cross-linking for 10 min at room temperature. Formaldehyde was quenched using glycine (0.125 M). Cross-linked cells were washed and pelleted twice with buffer A1, once with cell lysis buffer (140mM NaCl, 15mM Hepes, 1mM EDTA, 0.5mM EGTA, 1% Triton X100, 0.1% Sodium deoxycholate, 10mM sodium butyrate), followed by lysis in buffer containing, 140mM NaCl, 15mM Hepes, 1mM EDTA, 0.5mMEGTA, 1% Triton X100, 0.5% SDS, 0.5% N-Laurosylsarcosine, 0.1% sodium deoxycholate, 10mM sodium butyrate for 120 min at 4°C. Lysates were sonicated in Bioruptor sonicator to reach a fragment size window of 200-600 bp.

Chromatin was incubated overnight at 4°C with the following antibodies: for H3K9me3 ChIP using α-H3K9me3 (actif motif #39161, 3μg/IP) and for H3K4me3 using α-H3K4me3 (millipore #07-473, 3μg/IP) antibodies. The Magna ChIP A/G Chromatin Immunoprecipitation Kit (cat# 17-10085) was used following manufacturer’s instructions. Final DNA recovery was perfomed by classic phenol/chloroform DNA precipitation method using MAxtract high density tubes to maximize DNA recovery.

DNA fragments were then sequenced on an Illumina HiSeq 4000 apparatus, with paired-end 100 nt reads. Due to technical issues, only one replicate could be used for dsgoth31 input.

### ChIP-seq quality check: Validation of H3K4me3 enrichment around promoters and H3K9me3 on heterochromatic regions

Raw reads were trimmed using trim_galore (https://zenodo.org/record/5127899#.YbnMs73MLDc) with default parameters along with --paired, --clip_R1 9, --clip_R2 9, and --max_n 0. Mapping was performed using Bowtie2 (Langmead & Salzberg 2012) with --sensitive-local against the *D. melanogaster* r6.16 and *D. simulans* r2.02 genomes. Samtools was used to convert SAM to coordinated sorted BAM files, while sambamba (Tarasov et al. 2015) was used to filter for uniquely mapping reads and to remove duplicates (sambamba view -h -t 2 -f bam -F “[XS] == null and not unmapped and not duplicate”). For *D. melanogaster* datasets, we filtered available blacklisted regions (Amemiya et al. 2019) with bedtools. Finally coverage files containing reads per genome coverage (RPGC) were obtained with DeepTools (Ramírez et al. 2016) bamCoverage with --extendReads, --effectiveGenomeSize 129789873 for *D. melanogaster* available from the Deeptools suite, and --effectiveGenomeSize 121102921 computed with <u>unique-kmers.py</u> from khmer (https://github.com/dib-lab/khmer). Promoter regions were obtained with gencode_regions (https://github.com/saketkc/gencode_regions) and along with coverage files were used in DeepTools computeMatrix and plotProfile to build the average coverage of H3K4me3 and H3K9me3 around transcription start sites in both species and on chromosomes for H3K9me3. The corresponding profiles looked as expected (Supplemental Fig. S11, S12).

### ChIP-seq analysis

For each of the immunoprecipitated samples (H3K4me3, H3K9me3, input), TE read counts were computed at the family level using the TEcount module of TEtools (Lerat et al. 2017) and the list of TE sequences available at ftp://pbil.univ-lyon1.fr/pub/datasets/Roy2019/.

ChIP-seq counts were normalized across samples of the same species using the counts(normalize=T) function of DESeq2 1.26.0 (Love et al. 2014). This was done independently for each of the immunoprecipitated samples (H3K4me3, H3K9me3, input). We then performed a log-transformation using the rlogTransformation function of DESeq2, and subsequently considered mean values across replicates. We only kept genes expressed in ovaries. We chose to work on log-transformed values because log-transformation of count variables makes them fit normal assumption and thus makes them suitable for linear models. In addition, a ratio becomes a difference when log-transformed, which ensure the strict equivalence with the classical normalization approach consisting in dividing histone counts with input counts: log ([H3Kime3 counts] / [input counts]) = log(H3Kime3 counts) – log(input counts).

In order to quantify the associations between TE insertions and histone marks enrichment, we used the following linear models on log transformed read counts: histone mark (either H3K4me3 or H3K4me3) ∼ input + exon + intron + upstream + downstream, where “exon”, “intron”, “upstream”, and “downstream” are the numbers of TE insertions in exons, introns, 5 kb upstream sequences, and 5 kb downstream sequences, respectively. Size effects for each of these three factors were then recorded as the coefficients for the explanatory variables. To compute the contribution to total variance, we divided the Sum Square of the corresponding variables by the Total Sum Square, provided by the ANOVA of the linear model.

### Small RNA extraction, sequencing, and analyses

Small RNA extraction, sequencing, and analyses dedicated to TEs had already been performed and described in Mohamed *et al*. (Mohamed et al. 2020). Sequence files had been deposited in NCBI SRA under the accession number PRJNA644327.

Gene-derived small RNAs: Sequencing adapters were removed using cutadapt (Martin 2011), and 23-30 nt reads from one hand (considered as piRNAs) and 21 nt reads from the other hand (considered as siRNAs) were extracted using PRINSEQ lite (Schmieder & Edwards 2011), as described in (Mohamed et al. 2020). Reads were then aligned on previously masked genomes (see above, RNA-seq section) using bowtie --best (Langmead et al. 2009). Aligned reads were counted using eXpress (Roberts et al. 2011) and “tot_counts” were considered.

## Data Availability

The RNA-seq data generated in this study have been submitted to the NCBI BioProject database (https://www.ncbi.nlm.nih.gov/bioproject/) under accession number PRJNA795668. The ChIP-seq data generated in this study have been submitted to the NCBI BioProject database under accession number PRJNA796157. TE and gene annotations have been deposited to Zenodo doi: 10.5281/zenodo.7189887 (https://zenodo.org/record/7189887#.Y7RQ-KeZNH4).

Count tables for TE insertions, RNA-seq and ChIP-seq data are provided as Supplementary Files available at ftp://pbil.univ-lyon1.fr/pub/datasets/Fablet2023.

## Acknowledgments

We thank Francois Sabot, Matthieu Boulesteix, Vincent Mérel, and Daniel S Oliveira for useful discussions and technical help. We thank the GDR MobilET for useful discussions. We thank Gladys Mialdea, Justine Picarle, Sonia Janillon, and Nelly Burlet for technical help. This work was performed using the computing facilities of the CC LBBE/PRABI. Sequencing was performed by the GenomEast platform, a member of the “France Génomique” consortium (ANR-10-INBS-0009). This work was supported by Fondation pour la Recherche Médicale (grant DEP20131128536) and Agence Nationale de la Recherche (grant ExHyb ANR-14-CE19-0016-01). The authors declare no competing interests.

## Supporting Information files

**Supplemental_Fig_S1.** (*A*) Proportions of TE read counts in RNA-seq data relative to read counts corresponding to 1:1 orthologous genes and TEs. For each strain, two biological replicates are shown. (*B*) Number of TE insertions per functional region per strain, considering the 12,470 genes that are 1:1 orthologous between *D. melanogaster* and *D. simulans*. Upstream and downstream regions are 5 kb sequences directly flanking transcription units 5’ and 3’, respectively.

**Supplemental_Fig_S2.** (*A*) Positive correlations between per TE family RNA counts and family sequence abundance (in bp) (log10 transformed, Spearman correlations). (*B*) Positive correlations between per TE family RNA counts and TE-derived piRNA counts (log10 transformed, Spearman correlations).

**Supplemental_Table_S3.** Contribution to total Sum Squares in the models anova(lm(log10(RNA+1) ∼ log10(H3K4me3+1) + log10(H3K9me3+1) + log10(input+1))), calculated at the TE family level.

**Supplemental_Fig_S4. Detailed analysis of genes at the ends of distributions in Figure 3A**. (*A*) Analysis of mean expression difference, *D. melanogaster*, (*B*) *D. simulans*. (*C*) Analysis of mean H3K4me3 enrichment difference, *D. melanogaster*, (*D) D. simulans*. (*E*) Analysis of mean H3K9me3 enrichment difference, *D. melanogaster*, (*F) D. simulans*.

**Supplemental_Fig_S5. Analysis on 1:1 ortholog genes.** (*A*) Contribution of TE insertion numbers to gene expression total variance estimated using the linear model gene TPM (log) ∼ exon + intron + upstream + downstream, and (*B*) corresponding size effects. (*C*) Contribution of TE insertion numbers to gene H3K4me3 total variance estimated using the linear model gene H3K4me3 level (log) ∼ exon + intron + upstream + downstream, and (*D*) corresponding size effects. (*E*) Contribution of TE insertion numbers to gene H3K9me3 total variance estimated using the linear model gene H3K9me3 level (log) ∼ exon + intron + upstream + downstream, and (*F*) corresponding size effects. Colored bars: p-values < 0.05, empty bars: p-values > 0.05. Error bars are standard errors.

**Supplemental_Fig_S6. Separate analyses across TE classes** (A) Numbers of TE insertions per functional region per strain. Upstream and downstream regions are 5 kb sequences directly flanking transcription units 5’ and 3’, respectively. (B) Size effects to the contribution of TE insertion numbers to gene expression using the linear model gene TPM (log) ∼ exon + intron + upstream + downstream. (C) Size effects to the contribution of TE insertion numbers to gene H3K4me3 using the linear model gene H3K4me3 level (log) ∼ exon + intron + upstream + downstream. (D) Size effects to the contribution of TE insertion numbers to gene H3K9me3 using the linear model gene H3K9me3 level (log) ∼ exon + intron + upstream + downstream. Colored bars: p-values < 0.05, empty bars: p-values > 0.05. Error bars are standard errors.

**Supplemental_Fig_S7. Separate analyses across common and private TE insertions.** (A) Numbers of TE insertions per functional region per strain. Upstream and downstream regions are 5 kb sequences directly flanking transcription units 5’ and 3’, respectively. (B) Size effects to the contribution of TE insertion numbers to gene expression using the linear model gene TPM (log) ∼ exon + intron + upstream + downstream. (C) Size effects to the contribution of TE insertion numbers to gene H3K4me3 using the linear model gene H3K4me3 level (log) ∼ exon + intron + upstream + downstream. (D) Size effects to the contribution of TE insertion numbers to gene H3K9me3 using the linear model gene H3K9me3 level (log) ∼ exon + intron + upstream + downstream. Colored bars: p-values < 0.05, empty bars: p-values > 0.05. Error bars are standard errors.

**Supplemental_Fig_S8.** Correlation coefficients between gene-derived piRNAs and gene-derived 21 nt RNAs, between gene-derived piRNAs and gene H3K9me3 levels, and between gene-derived 21 nt RNAs and gene H3K9me3 levels. To the bottom are significance results for Wilcoxon rank tests comparing values for *D. melanogaster vs* values for *D. simulans*.

**Supplemental_Table_S9. Gene-derived piRNA production.**

From left to right : strain; 3^rd^ quartile of the distribution of gene-derived piRNA numbers; number of private TE-carrying genes; number of private TE-carrying genes with piRNA production higher than 3^rd^ quartile; number of private TE-carrying genes with piRNA production lower than private TE-carrying genes 3^rd^ quartile.

**Supplemental_Fig_S10. Validation of H3K4me3 enrichment around promoters.** Mean read coverage for H3K4me3 and H3K9me3 around Transcription start sites (TSS) of *D. melanogaster* and *D. simulans* datasets.

**Supplemental_Fig_S11. Validation of H3K9me3 enrichment on chromosomes.** Mean read coverage for H3K9me3 on chromosomes of *D. melanogaster* and *D. simulans* datasets. Validation of H3K9me3 enrichment in the heterochromatic chromosome 4 compared to other *D. melanogaster* and *D. simulans* chromosomes.

**Supplemental Files.** Available at ftp://pbil.univ-lyon1.fr/pub/datasets/Fablet2023 insertion_dnarep1_.txt

These files contain the numbers of insertions per gene per functional regions AFTER removing DNAREP1 insertions

table_NEWTPM_rna_dmel_modif.txt

table_NEWTPM_rna_dsim_modif.txt

These files contain the TPM obtained on genes from RNAseq data

table_NEWCOUNTS_rna_dmel_modif.txt

table_NEWCOUNTS_rna_dsim_modif.txt

These files contain the effective counts obtained on genes from RNAseq data

counts_chip_input_dmel_fbgn.txt

counts_chip_k4_dmel_fbgn.txt

counts_chip_k9_dmel_fbgn.txt

counts_chip_input_dsim_fbgn.txt

counts_chip_k4_dsim_fbgn.txt

counts_chip_k9_dsim_fbgn.txt

These files contain the counts obtained on genes from ChIPseq data

rna_te_dmel.txt

rna_te_dsim.txt

These files contain the counts obtained on TEs from RNAseq data

chip_input_et_dmel.txt

chip_k4_et_dmel.txt

chip_k9_et_dmel.txt

chip_input_et_dsim.txt

chip_k4_et_dsim.txt

chip_k9_et_dsim.txt

These files contain the counts obtained on TEs from ChIPseq data

