## Supplemental for "A quantitative, genome-wide analysis in *Drosophila* reveals transposable elements’ influence on gene expression is species-specific"

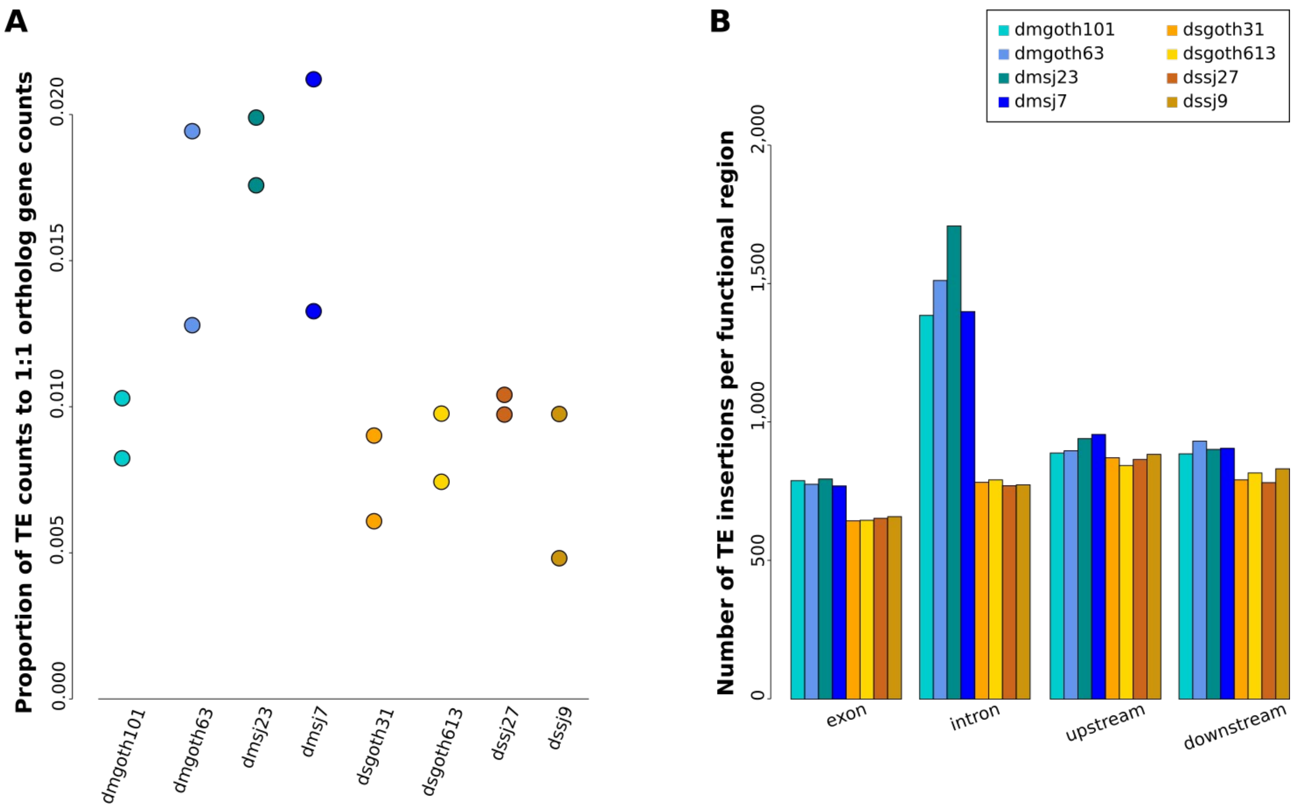


**Supplemental_Fig_S1**. (*A*) Proportions of TE read counts in RNA-seq data relative to read counts corresponding to 1:1 orthologous genes and TEs. For each strain, two biological replicates are shown. (*B*) Number of TE insertions per functional region per strain, considering the 12,470 genes that are 1:1 orthologous between *D. melanogaster* and *D. simulans*. Upstream and downstream regions are 5 kb sequences directly flanking transcription units 5′ and 3′, respectively.

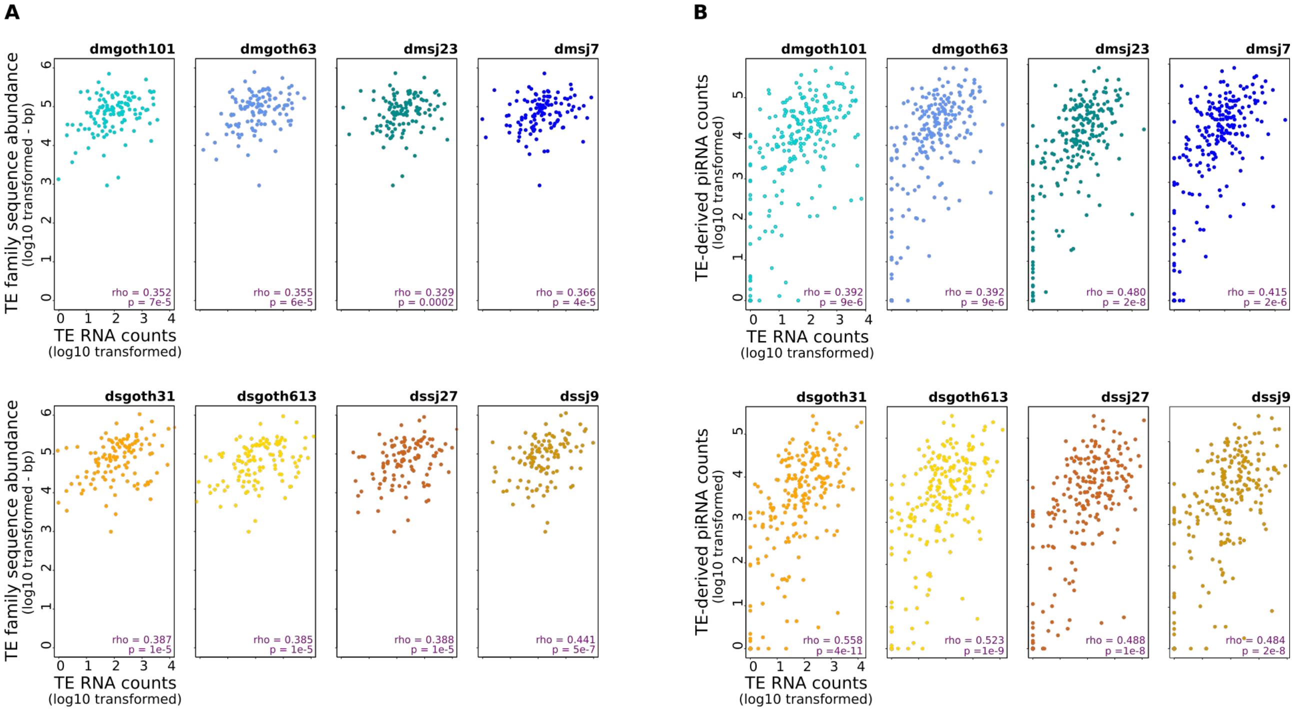


**Supplemental_Fig_S2.** (*A*) Positive correlations between per TE family RNA counts and family sequence abundance (in bp) (log10 transformed, Spearman correlations). (*B*) Positive correlations between per TE family RNA counts and TE-derived piRNA counts (log10 transformed, Spearman correlations).

**Supplemental_Table_S3**. Contribution to total Sum Squares in the models anova(lm(log10(RNA+1) ~ log10(H3K4me3+1) + log10(H3K9me3+1) + log10(input+1))), calculated at the TE family level.

| | | **H3K4me3** | **H3K9me3** | **input** |
|---|---|---|---|---|
| *D. melanogaster* | **dmgoth101** | 0.518 | 0.098 | 4e-5 |
| | **dmgoth63** | 0.407 | 0.052 | 0.049 |
| | **dmsj23** | 0.467 | 0.019 | 0.005 |
| | **dmsj7** | 0.514 | 0.126 | 0.006 |
| *D. simulans* | **dsgoth31** | 0.470 | 0.086 | 9e-4 |
| | **dsgoth613** | 0.488 | 0.063 | 0.002 |
| | **dssj27** | 0.483 | 0.118 | 0.006 |
| | **dssj9** | 0.408 | 0.060 | 2e-5 |

A

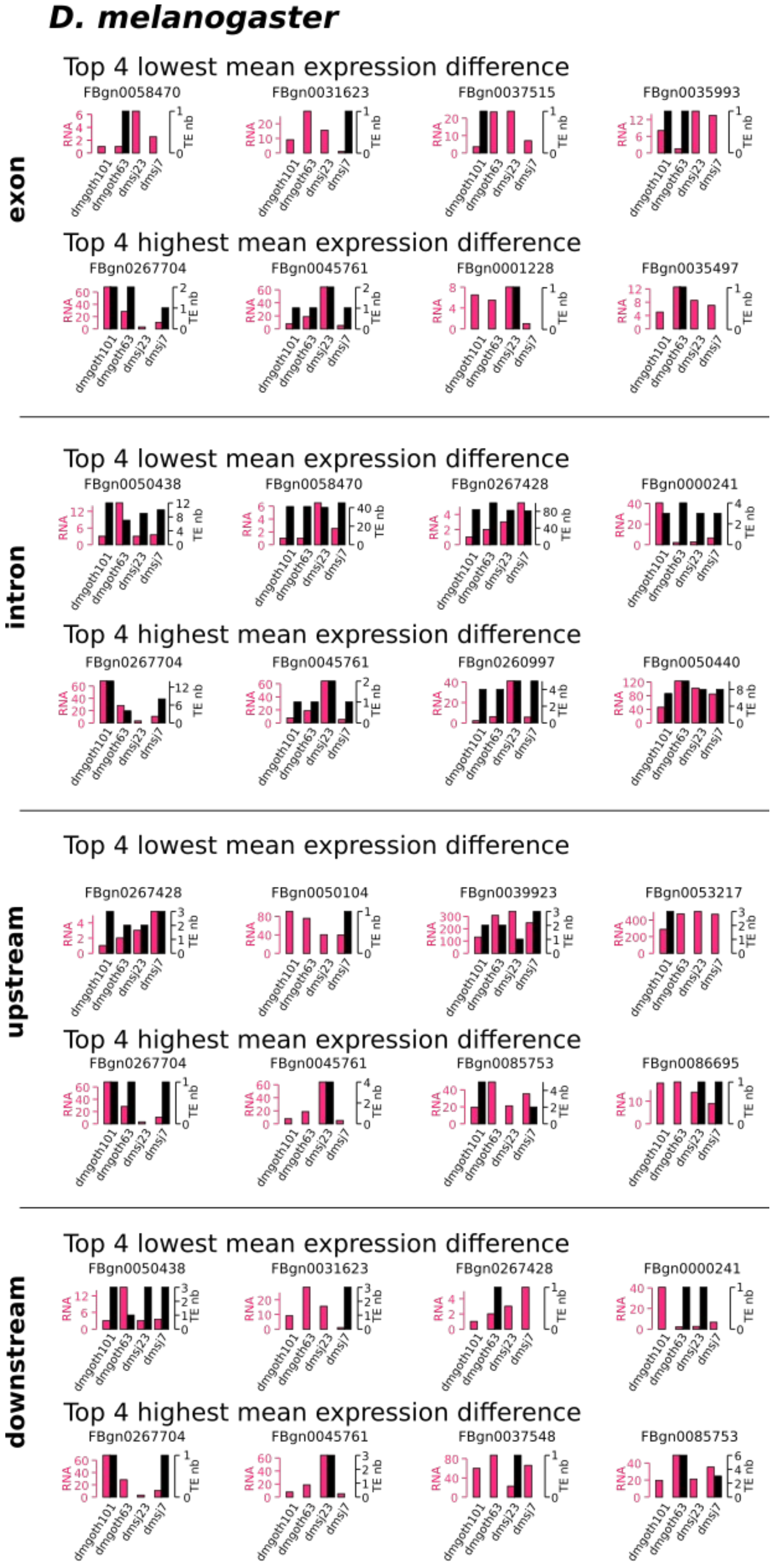
D. melanogaster
exon
intron
upstream
downstream
Top 4 lowest mean expression difference
Top 4 highest mean expression difference
FBgn0058470
FBgn0031623
FBgn0037515
FBgn0035993
FBgn0267704
FBgn0045761
FBgn0001228
FBgn0035497
FBgn0050438
FBgn0267428
FBgn0000241
FBgn0260997
FBgn0050440
FBgn0050104
FBgn0039923
FBgn0053217
FBgn0085753
FBgn0086695
FBgn0037548
RNA
TE nb
dmgoth101
dmgoth63
dmsj23
dmsj7

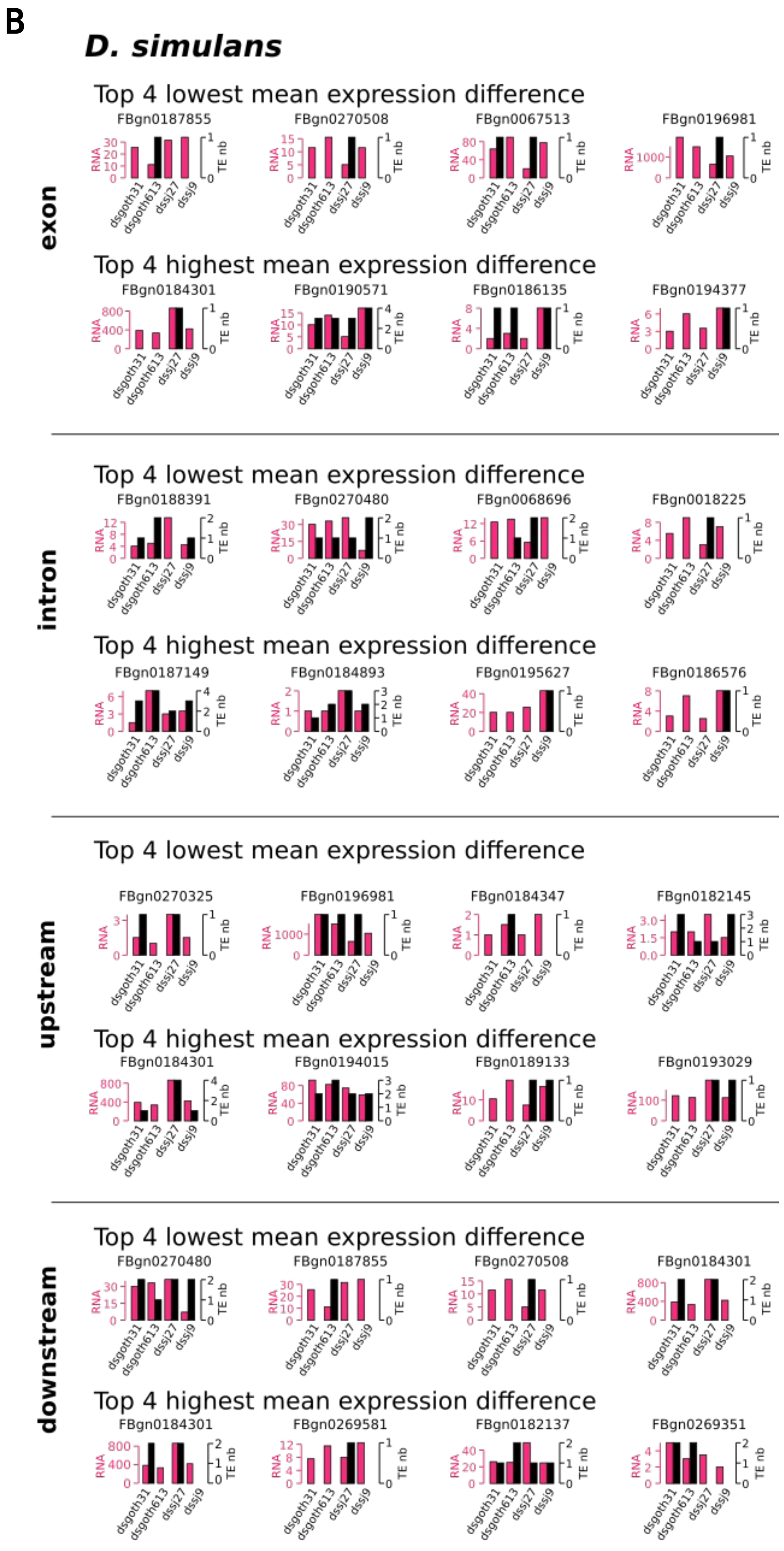
B
D. simulans
exon
Top 4 lowest mean expression difference
FBgn0187855
FBgn0270508
FBgn0067513
FBgn0196981
Top 4 highest mean expression difference
FBgn0184301
FBgn0190571
FBgn0186135
FBgn0194377
intron
Top 4 lowest mean expression difference
FBgn0188391
FBgn0270480
FBgn0068696
FBgn0018225
Top 4 highest mean expression difference
FBgn0187149
FBgn0184893
FBgn0195627
FBgn0186576
upstream
Top 4 lowest mean expression difference
FBgn0270325
FBgn0196981
FBgn0184347
FBgn0182145
Top 4 highest mean expression difference
FBgn0184301
FBgn0194015
FBgn0189133
FBgn0193029
downstream
Top 4 lowest mean expression difference
FBgn0270480
FBgn0187855
FBgn0270508
FBgn0184301
Top 4 highest mean expression difference
FBgn0184301
FBgn0269581
FBgn0182137
FBgn0269351
RNA
TE nb
dsgoth31
dsgoth613
dssj27
dssj9

C

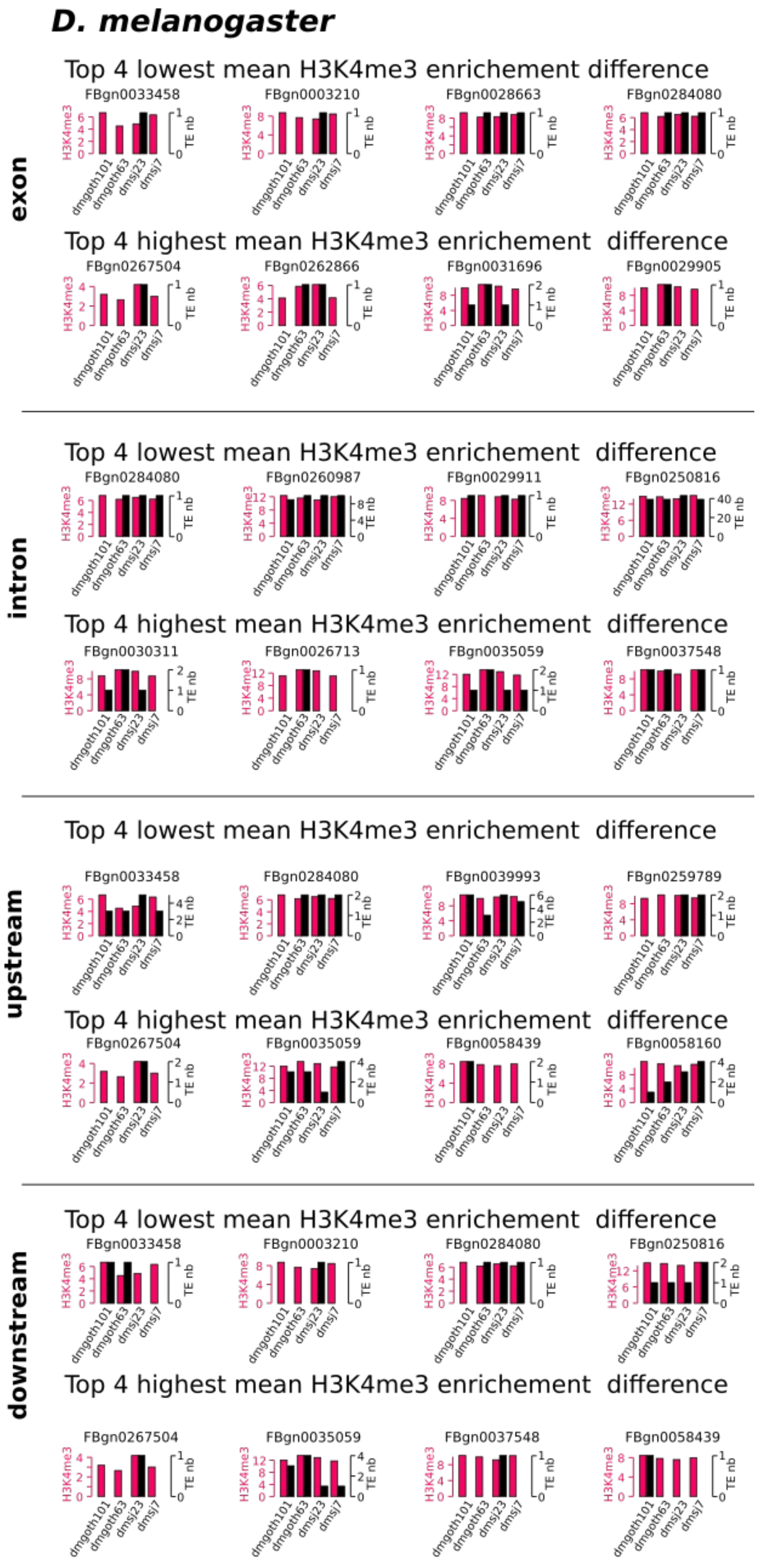
D. melanogaster
exon
Top 4 lowest mean H3K4me3 enrichement difference
FBgn0033458
FBgn0003210
FBgn0028663
FBgn0284080
Top 4 highest mean H3K4me3 enrichement difference
FBgn0267504
FBgn0262866
FBgn0031696
FBgn0029905
intron
Top 4 lowest mean H3K4me3 enrichement difference
FBgn0284080
FBgn0260987
FBgn0029911
FBgn0250816
Top 4 highest mean H3K4me3 enrichement difference
FBgn0030311
FBgn0026713
FBgn0035059
FBgn0037548
upstream
Top 4 lowest mean H3K4me3 enrichement difference
FBgn0033458
FBgn0284080
FBgn0039993
FBgn0259789
Top 4 highest mean H3K4me3 enrichement difference
FBgn0267504
FBgn0035059
FBgn0058439
FBgn0058160
downstream
Top 4 lowest mean H3K4me3 enrichement difference
FBgn0033458
FBgn0003210
FBgn0284080
FBgn0250816
Top 4 highest mean H3K4me3 enrichement difference
FBgn0267504
FBgn0035059
FBgn0037548
FBgn0058439
H3K4me3
TE nb
dmgoth101
dmgoth63
dmsj23
dmsj7

D

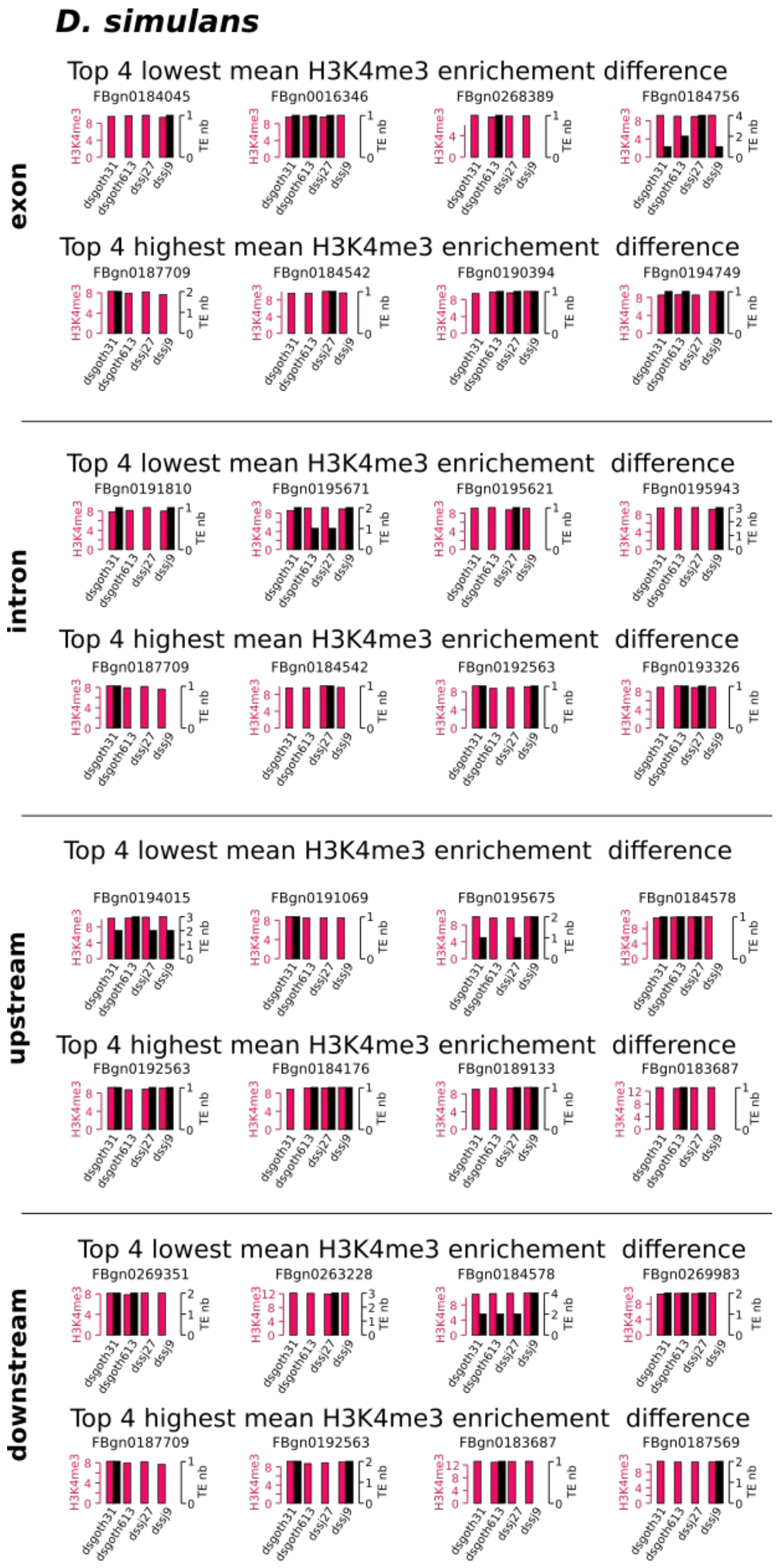
D. simulans
exon
Top 4 lowest mean H3K4me3 enrichement difference
FBgn0184045
FBgn0016346
FBgn0268389
FBgn0184756
Top 4 highest mean H3K4me3 enrichement difference
FBgn0187709
FBgn0184542
FBgn0190394
FBgn0194749
intron
Top 4 lowest mean H3K4me3 enrichement difference
FBgn0191810
FBgn0195671
FBgn0195621
FBgn0195943
Top 4 highest mean H3K4me3 enrichement difference
FBgn0187709
FBgn0184542
FBgn0192563
FBgn0193326
upstream
Top 4 lowest mean H3K4me3 enrichement difference
FBgn0194015
FBgn0191069
FBgn0195675
FBgn0184578
Top 4 highest mean H3K4me3 enrichement difference
FBgn0192563
FBgn0184176
FBgn0189133
FBgn0183687
downstream
Top 4 lowest mean H3K4me3 enrichement difference
FBgn0269351
FBgn0263228
FBgn0184578
FBgn0269983
Top 4 highest mean H3K4me3 enrichement difference
FBgn0187709
FBgn0192563
FBgn0183687
FBgn0187569
H3K4me3
TE nb
dsgoth31
dsgoth613
dssj27
dssj9

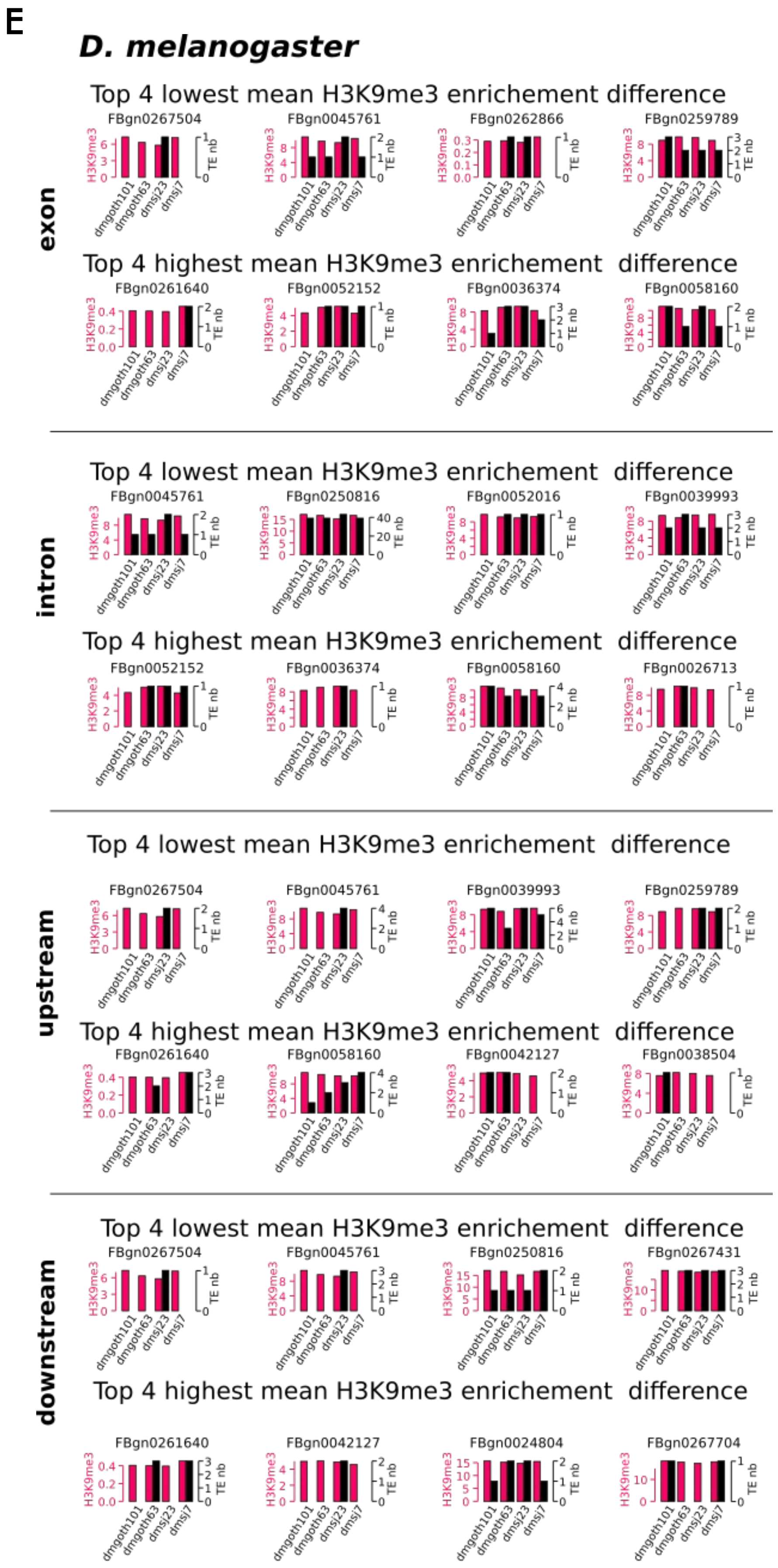
E
D. melanogaster
exon
Top 4 lowest mean H3K9me3 enrichement difference
FBgn0267504
FBgn0045761
FBgn0262866
FBgn0259789
Top 4 highest mean H3K9me3 enrichement difference
FBgn0261640
FBgn0052152
FBgn0036374
FBgn0058160
intron
Top 4 lowest mean H3K9me3 enrichement difference
FBgn0045761
FBgn0250816
FBgn0052016
FBgn0039993
Top 4 highest mean H3K9me3 enrichement difference
FBgn0052152
FBgn0036374
FBgn0058160
FBgn0026713
upstream
Top 4 lowest mean H3K9me3 enrichement difference
FBgn0267504
FBgn0045761
FBgn0039993
FBgn0259789
Top 4 highest mean H3K9me3 enrichement difference
FBgn0261640
FBgn0058160
FBgn0042127
FBgn0038504
downstream
Top 4 lowest mean H3K9me3 enrichement difference
FBgn0267504
FBgn0045761
FBgn0250816
FBgn0267431
Top 4 highest mean H3K9me3 enrichement difference
FBgn0261640
FBgn0042127
FBgn0024804
FBgn0267704
H3K9me3
TE nb
dmgoth101
dmgoth63
dmsj23
dmsj7

**F**

***D. simulans***

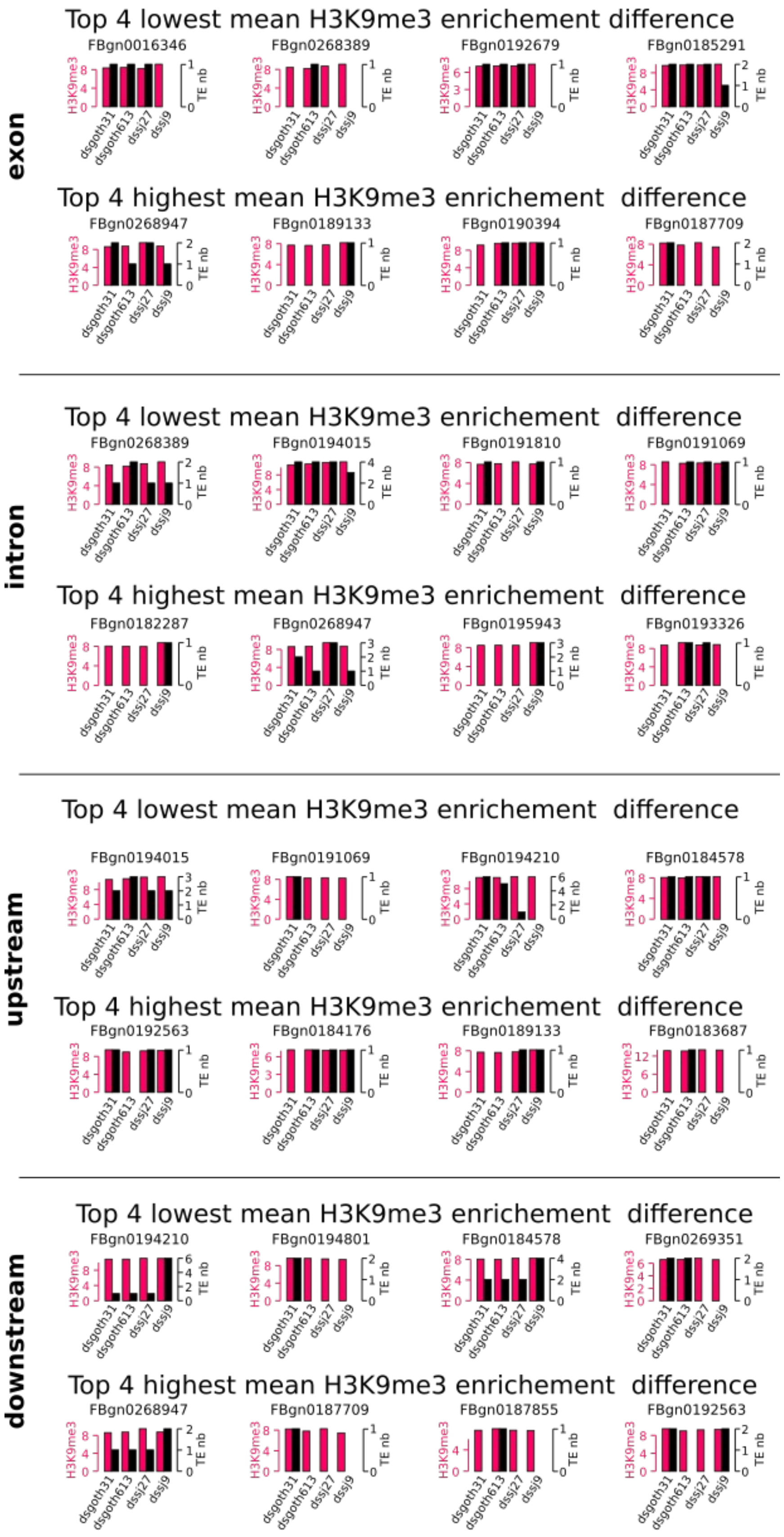


**Supplemental_Fig_S4. Detailed analysis of genes at the ends of distributions in Figure 3A.** (*A*) Analysis of mean expression difference, *D. melanogaster*, (*B*) *D. simulans*. (*C*) Analysis of mean H3K4me3 enrichment difference, *D. melanogaster*, (*D*) *D. simulans*. (*E*) Analysis of mean H3K9me3 enrichment difference, *D. melanogaster*, (*F*) *D. simulans*.

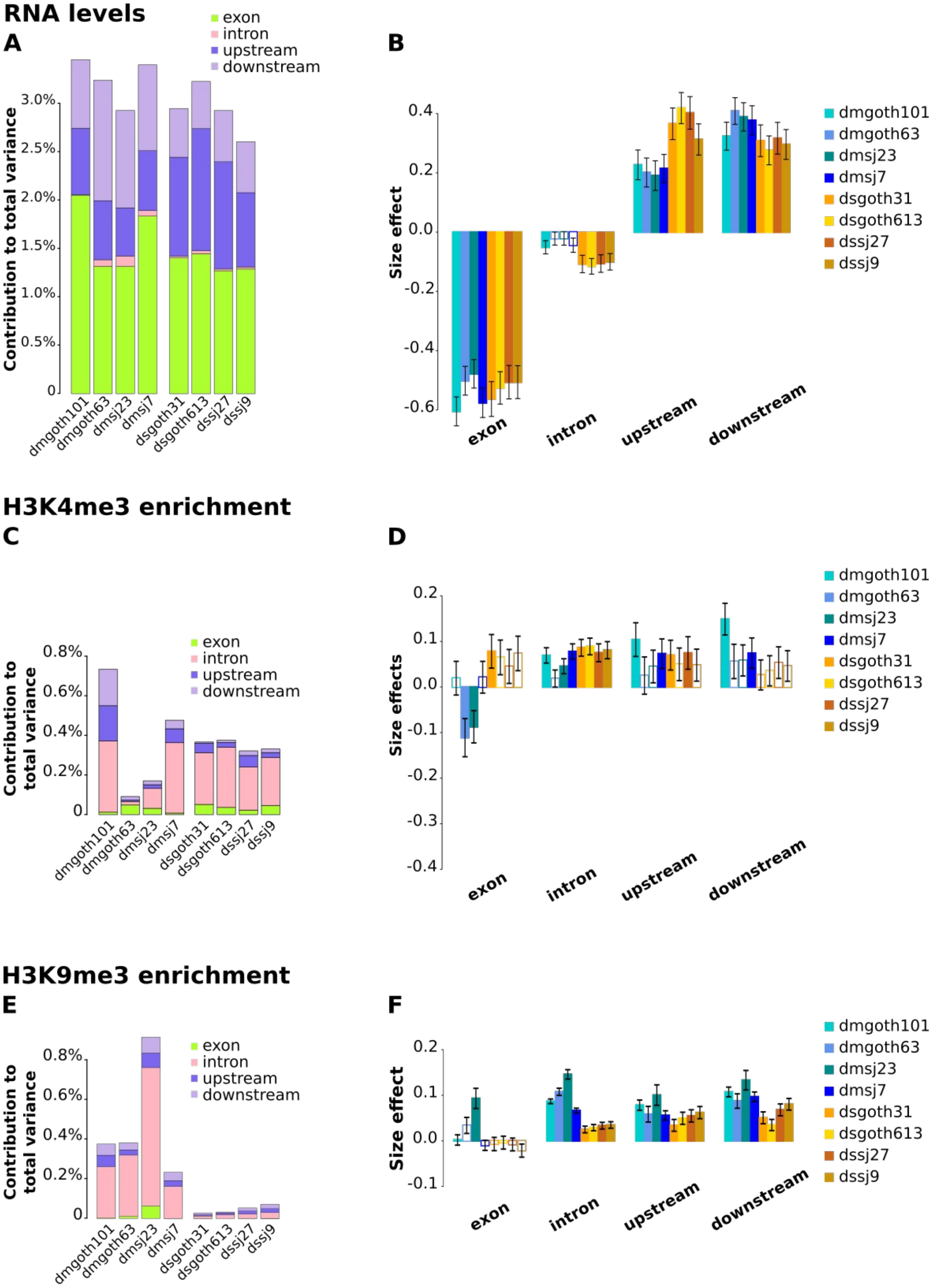


**Supplemental_Fig_S5. Analysis on 1:1 ortholog genes**. (*A*) Contribution of TE insertion numbers to gene expression total variance estimated using the linear model gene TPM (log) ~ exon + intron + upstream + downstream, and (*B*) corresponding size effects. (*C*) Contribution of TE insertion numbers to gene H3K4me3 total variance estimated using the linear model gene H3K4me3 level (log) ~ exon + intron + upstream + downstream, and (*D*) corresponding size effects. (*E*) Contribution of TE insertion numbers to gene H3K9me3 total variance estimated using the linear model gene H3K9me3 level (log) ~ exon + intron + upstream + downstream, and (*F*) corresponding size effects. Colored bars: p-values < 0.05, empty bars: p-values > 0.05. Error bars are standard errors.

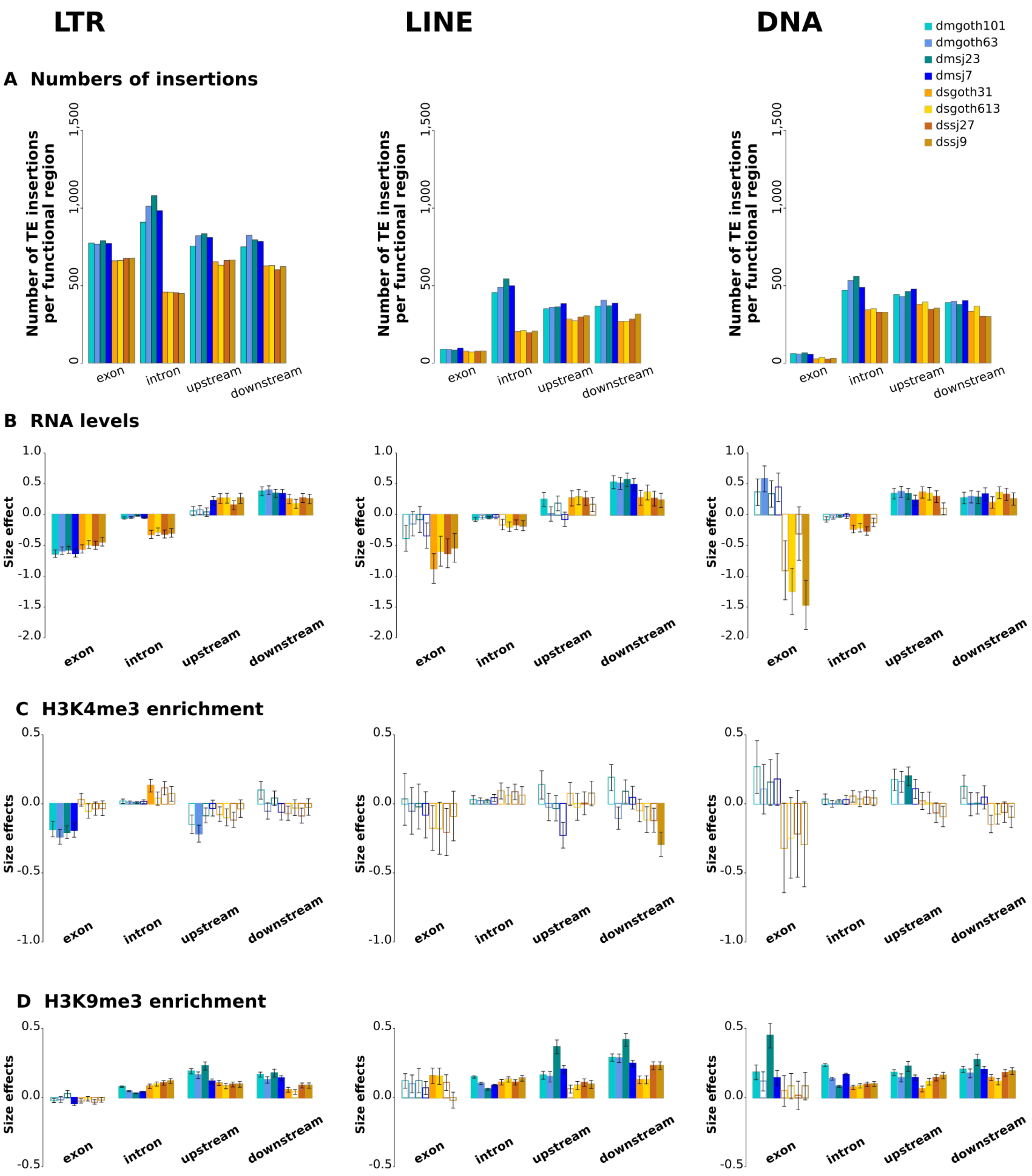


**Supplemental_Fig_S6. Separate analyses across TE classes** (A) Numbers of TE insertions per functional region per strain. Upstream and downstream regions are 5 kb sequences directly flanking transcription units 5' and 3', respectively. (B) Size effects to the contribution of TE insertion numbers to gene expression using the linear model gene TPM (log) ~ exon + intron + upstream + downstream. (C) Size effects to the contribution of TE insertion numbers to gene H3K4me3 using the linear model gene H3K4me3 level (log) ~ exon + intron + upstream + downstream. (D) Size effects to the contribution of TE insertion numbers to gene H3K9me3 using the linear model gene H3K9me3 level (log) ~ exon + intron + upstream + downstream. Colored bars: p-values < 0.05, empty bars: p-values > 0.05. Error bars are standard errors.

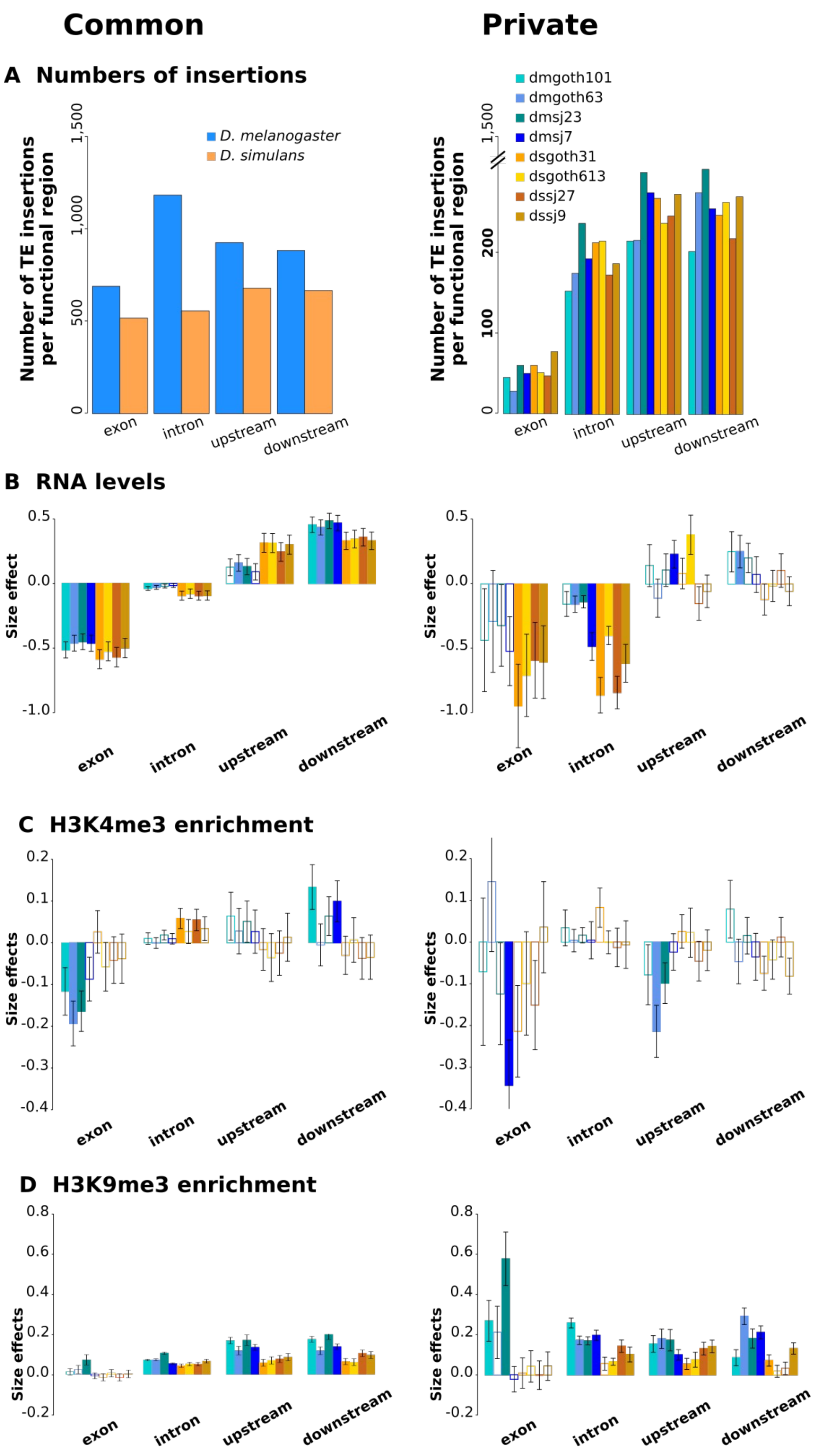


**Supplemental_Fig_S7. Separate analyses across common and private TE insertions.** (A) Numbers of TE insertions per functional region per strain. Upstream and downstream regions are 5 kb sequences directly flanking transcription units 5' and 3', respectively. (B) Size effects to the contribution of TE insertion numbers to gene expression using the linear model gene TPM (log) ~ exon + intron + upstream + downstream. (C) Size effects to the contribution of TE insertion numbers to gene H3K4me3 using the linear model gene H3K4me3 level (log) ~ exon + intron + upstream + downstream. (D) Size effects to the contribution of TE insertion numbers to gene H3K9me3 using the linear model gene H3K9me3 level (log) ~ exon + intron + upstream + downstream. Colored bars: p-values < 0.05, empty bars: p-values > 0.05. Error bars are standard errors.

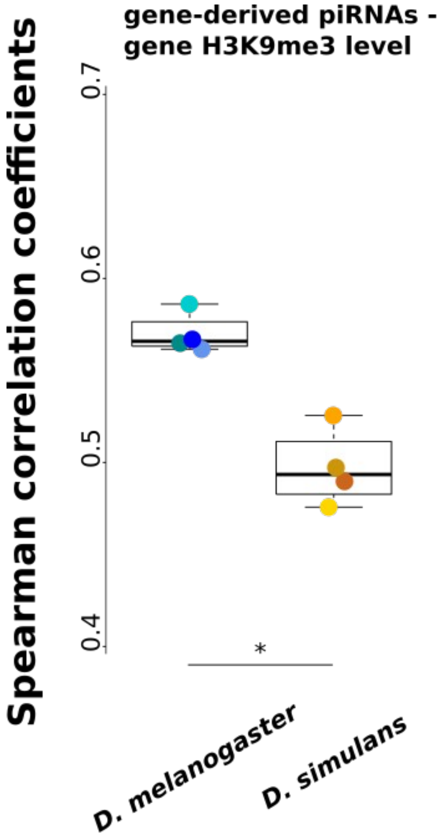


**Supplemental_Fig_S8.** Correlation cœfficients between gene-derived piRNAs and gene H3K9me3 levels,. To the bottom is significance result for Wilcoxon rank tests comparing values for *D. melanogaster vs* values for *D. simulans*.

**Supplemental_Table_S9. Gene-derived piRNA production.**

From left to right : strain; 3rd quartile of the distribution of gene-derived piRNA numbers; number of private TE-carrying genes; number of private TE-carrying genes with piRNA production higher than 3rd quartile; number of private TE-carrying genes with piRNA production lower than private TE-carrying genes 3rd quartile.

| | **3rd quartile (#piRNAs)** | **#genes with private TEs** | **>= 3rd qu** | **< 3rd qu** | **Chi-square** | **p-value** |
|---|---|---|---|---|---|---|
| **dmgoth101** | 60 | 29 | 14 | 15 | 8.3 | 0.0037 |
| **dmgoth63** | 68 | 34 | 21 | 13 | 24.5 | 7e-7 |
| **dmsj23** | 59 | 45 | 29 | 16 | 37.3 | 9e-10 |
| **dmsj7** | 43 | 47 | 40 | 7 | 90.5 | 1e-21 |
| **dsgoth31** | 23 | 48 | 32 | 16 | 44.4 | 2e-11 |
| **dsgoth613** | 17 | 52 | 30 | 22 | 29.6 | 5e-8 |
| **dssj27** | 15 | 58 | 33 | 25 | 31.4 | 2e-8 |
| **dssj9** | 14 | 57 | 28 | 29 | 17.6 | 2e-5 |

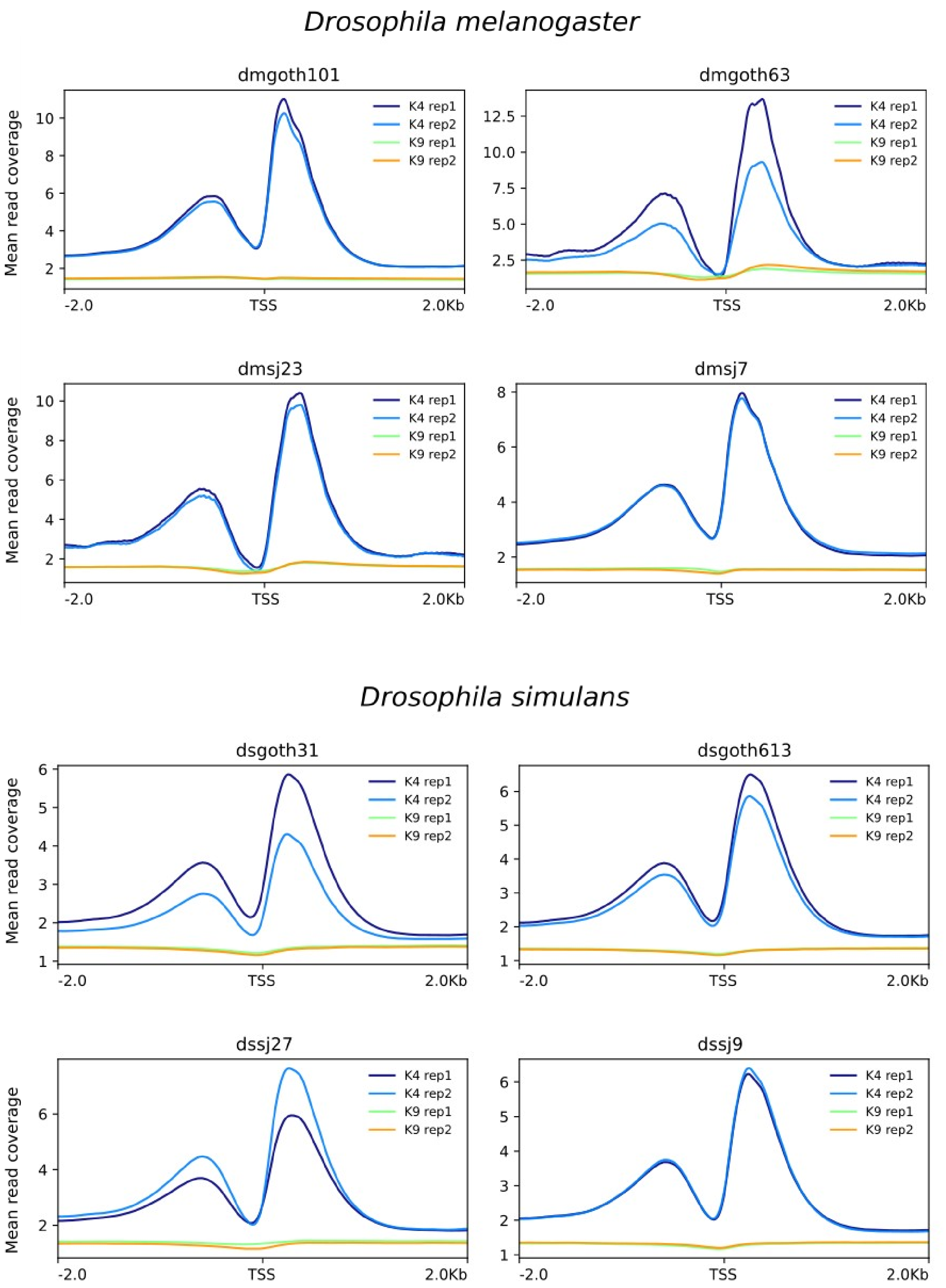


**Supplemental_Fig_S10. Validation of H3K4me3 enrichment around promoters.**
Mean read coverage for H3K4me3 and H3K9me3 around Transcription start sites (TSS) of *D. melanogaster* and *D. simulans* datasets.

*D. melanogaster*

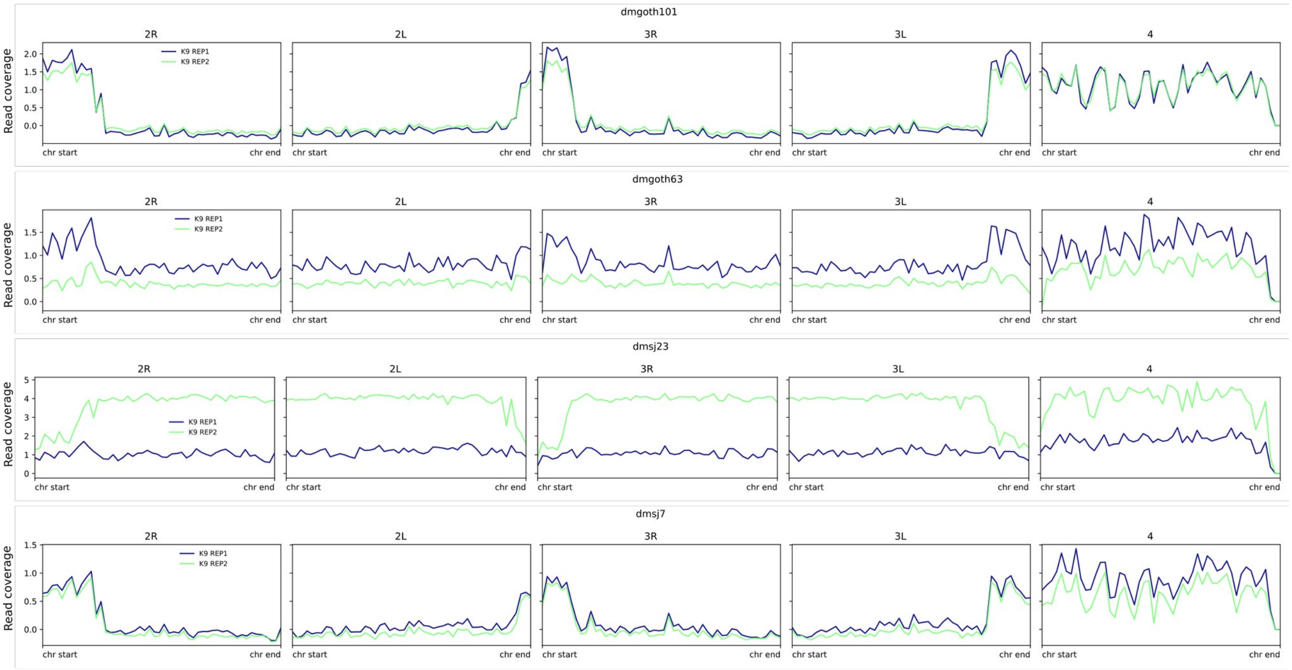


*D. simulans*

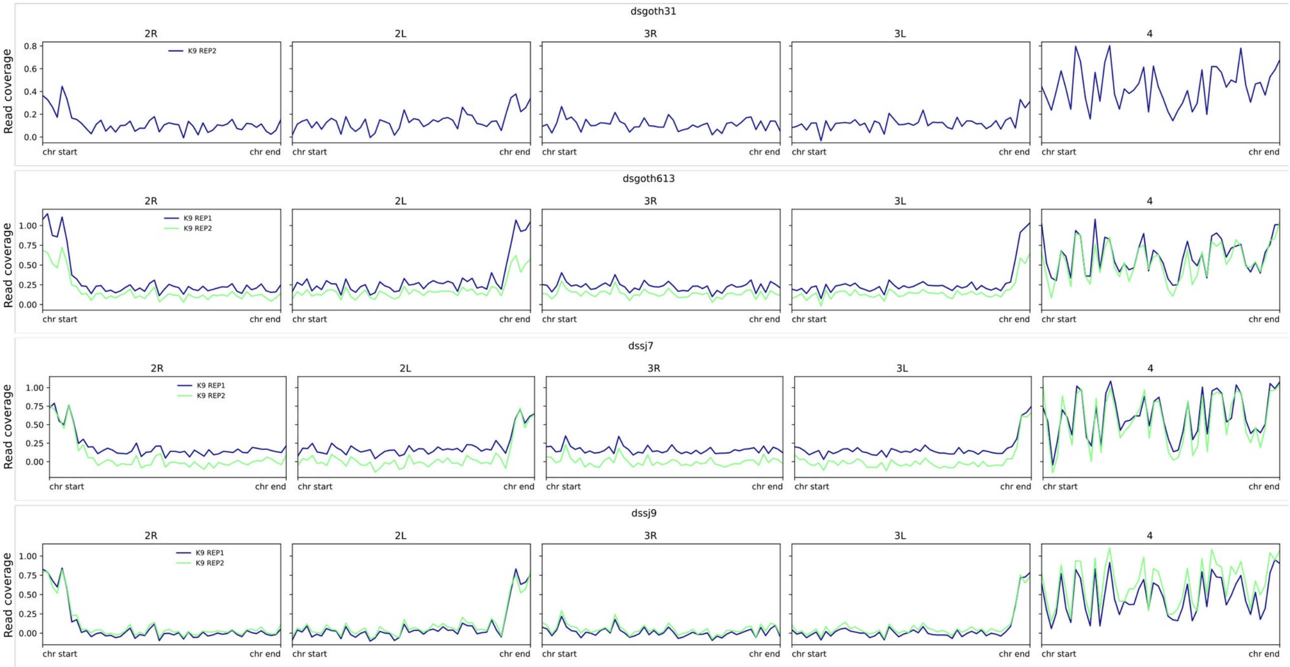


**Supplemental_Fig_S11. Validation of H3K9me3 enrichment on chromosomes.**
Mean read coverage for H3K9me3 on chromosomes of *D. melanogaster* and *D. simulans* datasets. Validation of H3K9me3 enrichment in the heterochromatic chromosome 4 compared to other *D. melanogaster* and *D. simulans* chromosomes.
